# Human Saposin B Ligand Binding and Presentation to α-Galactosidase A

**DOI:** 10.1101/2024.04.04.584535

**Authors:** Thomas K. Sawyer, Efecan Aral, James V. Staros, Cedric E. Bobst, Scott C. Garman

## Abstract

Sphingolipid activator protein B (saposin B; SapB) is an essential activator of globotriaosylceramide (Gb3) catabolism by α-galactosidase A. However, the manner by which SapB stimulates α-galactosidase A activity remains unknown. To uncover the molecular mechanism of SapB presenting Gb3 to α-galactosidase A, we subjected the fluorescent substrate globotriaosylceramide-nitrobenzoxidazole (Gb3-NBD) to a series of biochemical and structural assays involving SapB. First, we showed that SapB stably binds Gb3-NBD using a fluorescence equilibrium binding assay, isolates Gb3-NBD from micelles, and facilitates α-galactosidase A cleavage of Gb3-NBD *in vitro*. Second, we crystallized SapB in the presence of Gb3-NBD and validated the ligand-bound assembly. Third, we captured transient interactions between SapB and α-galactosidase A by chemical cross-linking. Finally, we determined the crystal structure of SapB bound to α-galactosidase A. These findings establish general principles for molecular recognition in saposin:hydrolase complexes and highlight the utility of NBD reporter lipids in saposin biochemistry and structural biology.

## INTRODUCTION

Sphingolipid activator proteins (saposins, Saps) are non-enzymatic glycoproteins that bind sphingolipid cargoes and present them to lysosomal hydrolases^1^. Four mature saposins (SapA, SapB, SapC, and SapD) result from proteolysis of the precursor prosaposin as it traffics to the lysosome^2,3^. Each of the four mature saposins has preferred cargoes and presents those cargoes to different hydrolases in the lysosome. The saposin family members are disulfide-linked, four-helix bundles that usually form dimers^4^.

Consistent with their role in solubilizing sphingolipids, saposins are dynamic molecules. The various crystal structures of the lysosomal saposins reveal large conformational and quaternary structural changes that can be induced by the presence of lipophilic cargo depending on pH and temperature^5–7^.

Unlike the other saposins, SapB appears as an obligate dimer in solution^8^. Moreover, it is capable of binding sphingolipids and phospholipids at both neutral and lysosomal pH^9,10^. Unsurprisingly, the first crystal structure of SapB showed two V-shaped protein monomers forming an asymmetric dimer with some density for an *E. coli* membrane-extracted phospholipid that copurified with the protein^11^.

SapB presents a variety of ligands to several lysosomal hydrolases, including arylsulfatase A, β-galactosidase (GLB1), and α-galactosidase A (GLA)^12^. After presentation of the ligand to the hydrolase, the lysosomal enzyme cleaves the cargo. One cargo, globotriaosylceramide (Gb3), is presented exclusively by SapB to GLA for removal of the terminal α-galactose, and failure of this process leads to accumulation of Gb3 in the lysosome and ultimately results in Fabry disease^13^. Other defects in SapB presentation of cargo lead to metachromatic leukodystrophy due to the accumulation of substrates of arylsulfatase A^14^.

These observations raise the question of how saposins activate lysosomal hydrolases. One example of a mechanism of activation derives from the crystal structure of SapA bound to the lysosomal hydrolase β-galactosylceramidase (GALC)^15^. The structure showed a 2:2 SapA:GALC hetero-tetramer with open hydrophobic channels connecting the binding cavity of the SapA dimer to the active sites of both GALC proteins. The binary complex formed in the presence of lauryldimethylamine-*N*-oxide detergent (LDAO), a ligand that interacted with an open conformation of SapA in a ‘lipoprotein disc’ structure^16^. To date, a ternary complex of saposin, sphingolipid, and enzyme remains to be detected by any method *in vitro*.

To address the molecular mechanism of cargo presentation by SapB, we studied the interactions of SapB and GLA with the ligand Gb3-nitrobenzoxidazole (Gb3-NBD), a fluorescent derivative of the physiological substrate Gb3. First, we show that SapB binds to Gb3-NBD, resulting in a soluble lipoprotein complex that stimulates GLA activity. Second, we determined a crystal structure of SapB grown in the presence of Gb3-NBD and performed in-solution molecular dynamics simulations to affirm its ligand-bound state. Third, using covalent cross-linking to trap the transient saposin-hydrolase complex, we show that SapB makes a direct, ligand-dependent interaction with GLA. Finally, we determined the cocrystal structure of SapB bound to GLA and use in-solution molecular dynamics simulations to interpret the structure. These findings present a *modus operandi* for studying other saposin-ligand and saposin-hydrolase interactions.

## RESULTS

### SapB binds and solubilizes Gb3-NBD in a water-soluble lipoprotein complex

Although weakly fluorescent in water, the fluorescent reporter nitrobenzoxidazole (NBD) strongly fluoresces in hydrophobic environments^17–21^. When attached to the acyl chain of lipids, the NBD fluorescence quenches when the fluorophore reaches a lipid-water interface^22–24^. The critical micelle concentrations of Gb3 and acyl-chain-labeled NBD-phospholipids are nanomolar to low micromolar^25,26^. Importantly, Gb3-NBD behaves similarly to endogenous Gb3 substrate in Fabry disease cell models^27^. The addition of exogenous Gb3-NBD to cells containing the enzyme sphingolipid ceramide *N-*deacylase leads to the conversion of Gb3-NBD to a deacylated form, lyso-Gb3^28^. Moreover, recombinant GLA cleaves the terminal α-galactose from Gb3-NBD, generating the product lactosylceramide-NBD, *in vitro*^29^. Thus, the substrate analog Gb3-NBD can be a useful reporter for SapB binding to Gb3.

Saposins bind lipids within their hydrophobic interior pockets^30^. We hypothesized that binding of SapB to Gb3-NBD would result in a fluorescence increase of the NBD group, indicative of acyl chain encapsulation in the hydrophobic pocket of the SapB dimer (Fig. 1a). By titrating SapB against a fixed amount of Gb3-NBD, a large fluorescence increase occurred, generating a sigmoidal curve. We used lysozyme as a negative control and purified recombinant SapD as a saposin specificity control. After a one-hour incubation at pH 6.5, we observed that SapB bound Gb3-NBD with a 5.8 ± 0.5 μM apparent dissociation constant (Kd_app_). The complex was stable for twenty-one days, with a measured 7.3 ± 0.5 μM Kd_app_ (Fig. 1b). There was no fluorescence change with the lysozyme control and low but detectable change with the SapD control. To test for the exposure of the fluorophore to the aqueous media, we added an excess of the fluorescent quencher sodium iodide (NaI)^31–33^. NaI had minimal effect on the Gb3-NBD fluorescence, suggesting that the NBD was in the binding cavity of SapB and inaccessible to the iodide (Fig. 1b). We saw similar fluorescence curves at pH 6.5 and 4.6, the pH of the lysosome (Supplementary Fig. 1a). SapD titration against Gb3-NBD resulted in a sigmoidal fluorescence change only at acidic pH (38 ± 2 μM Kd_app_), with lower maximum binding than SapB.

**Figure 1:**
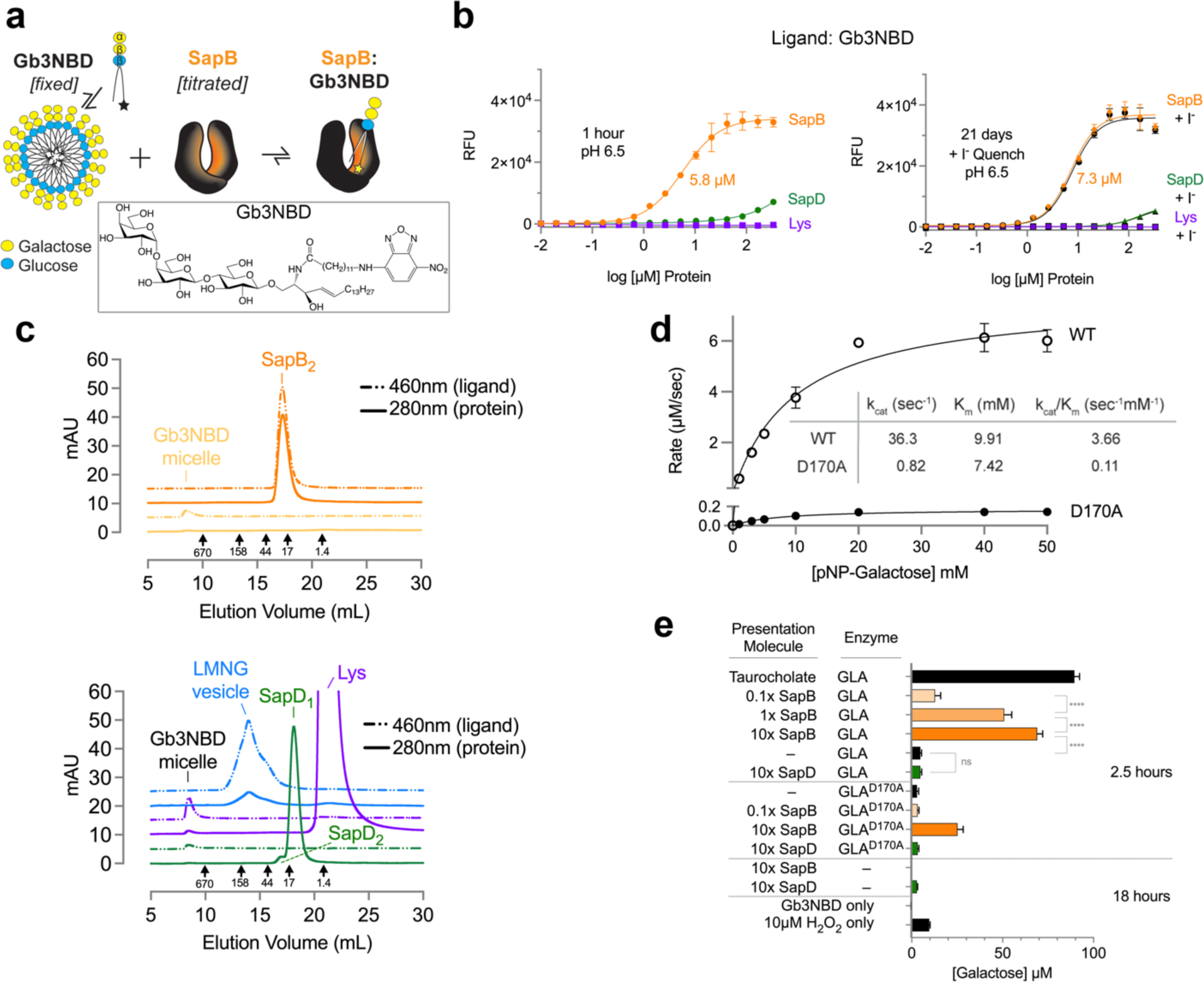
SapB binds, extracts, and presents Gb3 to GLA for catabolism. **a** Schematic of the Gb3-NBD equilibrium binding assay, in which a serial concentration of SapB is titrated against a fixed micellar concentration of Gb3-NBD. Due to the chemical property of the NBD moiety on the C12 fatty acyl chain, increased fluorescent emission is expected in a hydrophobic environment, such as the interior cavity of SapB (yellow star). The chemical structure of Gb3-NBD is boxed. **b** *Left*: The equilibrium binding of SapB (orange, circle), the saposin specificity control SapD (green, triangle), and negative control lysozyme (Lys, purple, square) to Gb3-NBD after one hour at pH 6.5. The apparent dissociation constants are labeled. *Right*: The same assay was analyzed twenty-one days later and then after the addition of iodide, a hydrophilic NBD-quencher. **c** *Top*: Comparison of the size-exclusion chromatograms of SapB:Gb3-NBD (orange) and Gb3-NBD micelles (yellow), in which dual wavelength monitoring tracks both protein (280nm, solid line) and Gb3-NBD ligand (460nm, dotted line) absorbance. *Bottom*: The corresponding chromatograms of the saposin specificity control SapD:Gb3-NBD (green), the negative control lysozyme:Gb3-NBD (purple), and the positive control, lipid-solubilizing detergent LMNG:Gb3-NBD (blue). **d** The kinetics of wild type GLA versus the catalytic nucleophile mutant GLA^D170A^. **e** The hydrolysis of Gb3-NBD, measured by an increase in the concentration of free galactose, after the time-specified incubation with the activator proteins SapB (orange) and SapD (green) with either wild type GLA or GLA^D170A^. The positive control sodium taurocholate, assay control H_2_O_2_, and other negative controls are shown in black. The error bars in panels represent the standard deviation between replicates.

As a substrate-specificity control, we assessed saposin binding to NBD-labeled ceramide (Cer-NBD), which lacks the carbohydrate head group of Gb3-NBD. As expected, SapB bound Cer-NBD with a much weaker affinity at both pH 6.5 (> 95 ± 9 μM Kd_app_) and pH 4.6 (> 129 ± 20 μM Kd_app_) at 21 days (Supplementary Fig. 1b,c). Overall, SapB has an approximate 17-fold greater affinity for Gb3-NBD compared to Cer-NBD, consistent with SapB presenting ligands to hydrolases that catabolize galactose-containing glycosphingolipids. The selectivity of SapB for Gb3 versus Cer was previously observed in a comprehensive competitive lipid binding study^34^.

In contrast to SapB, SapD binding to Cer-NBD was undetectable at either pH, suggesting that the NBD moiety may not be buried in the hydrophobic interior pocket of SapD or that SapD does not directly bind Cer-NBD (Supplementary Fig. 1b,c). Given our results, it is not clear how SapD specifically enhances ceramide degradation *in vitro* and in cells^35,36^.

To isolate complexes of SapB and Gb3-NBD, we performed size-exclusion chromatography (SEC) at pH 6.5. Simultaneous monitoring of the protein at 280nm and Gb3-NBD at 460nm allowed us to separate SapB:GB3-NBD complex from micelles. Absent a carrier molecule, there was a significant loss of Gb3-NBD to the column relative to the input, but we detected a small amount of Gb3-NBD micelles eluting at the column void volume. However, when mixed with SapB, there was a significant recovery of Gb3-NBD that comigrated with SapB, away from the micellar void elution (Fig. 1c). Altogether, the results show that SapB extracts Gb3-NBD from its micelle and forms a water-soluble lipoprotein complex. In comparison, the negative control lysozyme and the saposin specificity control SapD did not extract Gb3-NBD from the micelles (Fig. 1c). The delayed migration of the lysozyme control on the sizing column suggests that it interacts with the dextran-agarose column matrix^37^. We further employed the lipid-solubilizing detergent lauryl maltose neopentyl glycol (LMNG) as a positive control to solubilize Gb3-NBD. LMNG solubilized Gb3-NBD, shifting it out of the void volume and into a known LMNG elution profile on the column^38^ (Fig. 1c). In the presence and absence of Gb3-NBD, SapB and the other proteins ran at similar elution volumes on the sizing column (Supplementary Fig. 1d). Overall, the differences in Gb3-NBD absorbance between the samples reflect the ability of the carrier molecule to bind and solubilize the lipid.

### SapB activates GLA catabolism of Gb3-NBD

To demonstrate the activation of GLA by SapB, we performed enzyme kinetics. We expressed and purified from insect cells both wild-type GLA and GLA^D170A^, in which a catalytic nucleophile is removed^39^. Michaelis-Menten kinetic analysis at pH 5 using the synthetic substrate *p-*nitrophenyl (*p*NP)-galactose revealed that GLA^D170A^ (K_m_ 7.4 mM; k_cat_ 0.82 s^-1^) had about 2.3% the maximal activity of wild type (K_m_ of 9.9 mM; k_cat_ of 36.3 s^-1^) (Fig. 1d).

With the wild-type and D170A variant GLA enzymes, we also assessed the catabolism of Gb3-NBD in an enzyme-activator assay with SapB and SapD as presentation molecules. We detected the release of free galactose from Gb3-NBD in the presence of SapB in a concentration-dependent manner after a 2.5-hour incubation in an acidic buffer (Fig. 1e). At a molar equivalence with GLA, SapB stimulated the release of ∼50 μM galactose from the ∼200 μM input Gb3-NBD, which was slightly increased by raising SapB to a tenfold molar excess. Overall, SapB stimulated galactose cleavage from Gb3-NBD like the detergent taurocholate. In contrast, introducing SapD at a tenfold molar excess to GLA resulted in the release of ∼5 μM galactose, a detection two-fold above SapD alone control levels and slightly above the minimum of the assay. With a tenfold molar excess of SapB present, GLA^D170A^ cleaved ∼25 μM of the total Gb3-NBD, consistent with the loss of a catalytic residue (Fig. 1e). In these experiments, we showed the specificity of SapB in activating GLA catabolism of Gb3-NBD.

### Crystals of SapB in the presence of Gb3-NBD

To determine the molecular basis for SapB binding and presenting Gb3, we expressed and purified recombinant human SapB from *E. coli* and co-crystallized it at lysosomal pH in the presence of Gb3-NBD. Crystals took months to grow and were bright yellow, strongly indicative of NBD from the ligand in the crystal (Fig. 2a). We performed multiple experiments to test for the formation of the SapB:Gb3-NBD complex in the crystals. First, several dozen crystals were individually harvested and washed, and they retained their bright yellow color. Second, we dissolved washed crystals in PBS pH 6.5 and measured the NBD fluorescence emission spectrum, leading to a peak 472.5-fold above the crystal buffer control (Fig. 2a). Third, we dissolved washed crystals in 0.1 % formic acid, subjected the sample to liquid-chromatography mass spectrometry (LC-MS) analysis, which indicated the presence of both SapB and Gb3-NBD (Fig. 2b).

**Figure 2:**
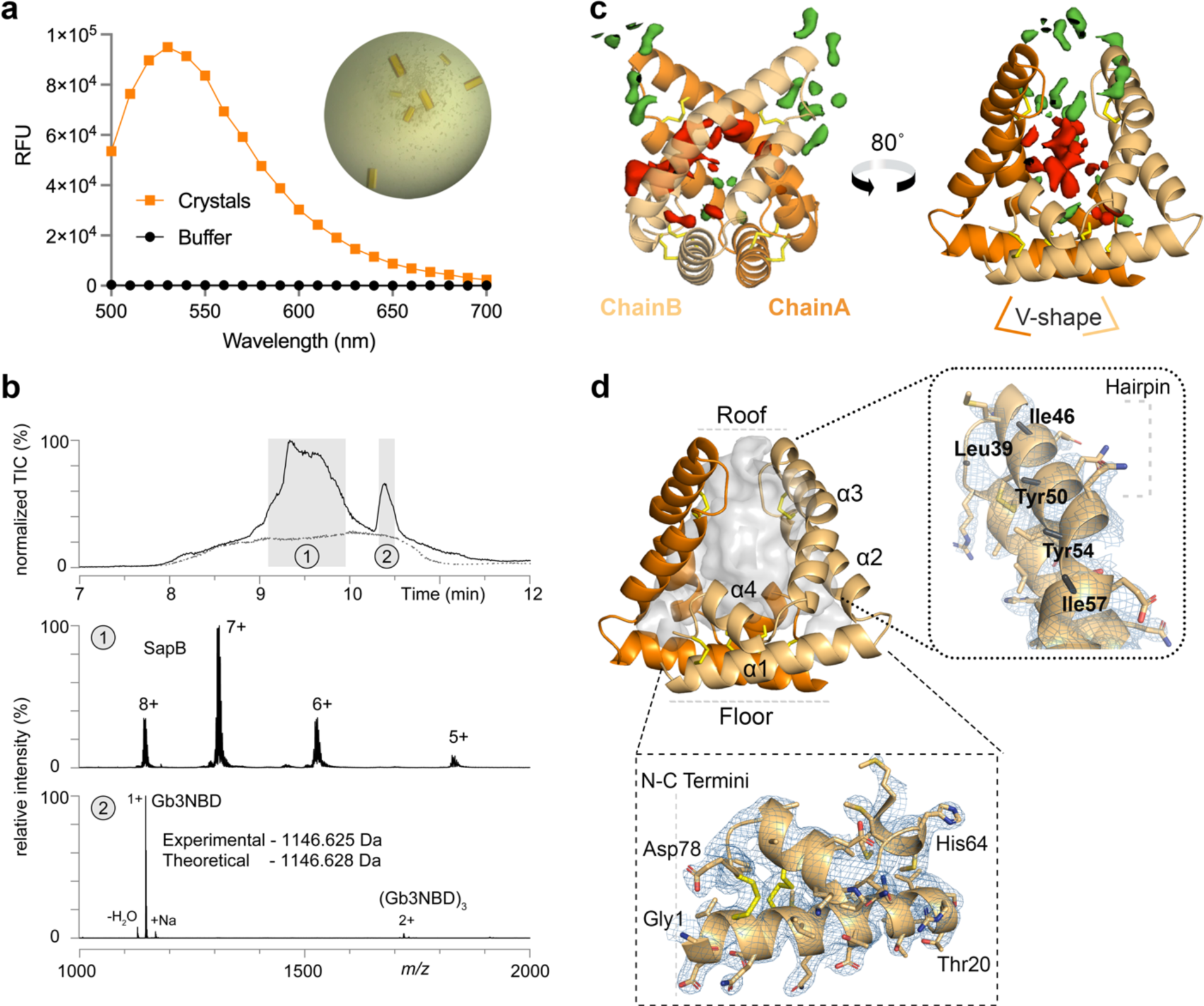
Co-crystals of SapB and Gb3-NBD show the presence of ligand. **a** The NBD fluorescence emission spectrum of several dozen SapB:Gb3-NBD crystals, individually looped and twice-washed, measured in a plate reader against a crystal buffer control. An optical image of the SapB:Gb3-NBD crystals shows the intense yellow hue of the crystals. **b** LC-MS analysis of the SapB:Gb3-NBD crystals. *Top*: Two distinct peaks were observed in the total ion chromatograms (TIC) for the crystal sample vs blank. Mass spectra produced within the shaded elution time windows (1 & 2) identify 1 as SapB (middle) and 2 as Gb3-NBD (bottom) with charge states labeled. **C** The structure of the SapB dimer, formed by two nearly symmetric V-shaped monomer chains with disulfide bonds indicated (yellow sticks). The mFo-DFc (green–positive/ red–negative at 2.5 σ) near chain A shows disjointed positive density surrounding the pocket, suggestive of a disordered Gb3 headgroup, and negative density in the interior hydrophobic pocket, consistent with bulk solvent rendering of disordered acyl chains. **d** The structure of the SapB dimer with its open cavity displayed (grey) and helices labeled. *Top*: The hairpin of chain B has several non-polar residues lacking 2mFo-DFc (blue, 1.0 σ) density beyond the beta carbon (black sticks). *Bottom*: The N-C terminal helical floor has unambiguous 2mFo-DFc (blue, 1.0 σ) density for all residues. Landmark residues for orientation are indicated.

The SapB structure revealed a homodimer consisting of helical V-shaped monomers that form a ∼2,900 Å^3^ binding cavity lined with hydrophobic residues. Given the above evidence for Gb3-NBD in the crystals, we were surprised to see only weak density in the ligand-binding cavity (Fig. 2c). Additionally, several hydrophobic residues on one face of a helix in the binding cavity showed little density beyond the beta carbon (Fig. 2d). We interpret the result as a collection of disordered ligands in the hydrophobic cavities of the SapB molecules in the crystalline state.

### Collapse of the hydrophobic binding pocket in the absence of lipid cargo

Because the SapB binding pocket contained many exposed hydrophobic residues in the SapB:Gb3-NBD structure, we used molecular dynamics (MD) simulations to investigate hydrophobic ligand binding in the SapB dimer. We used the coordinates from the X-ray structure as a starting point for MD simulations. Starting with the SapB:Gb3-NBD crystal structure, we modeled one Gb3-NBD ligand and then a non-overlapping, symmetric pair of ligands into the interior pocket. We performed MD simulations on the zero-, one- and two-Gb3-NBD occupied SapB dimers and looked at changes in the protein chain and dimer volume during the simulations (Fig. 3a). During 2 μs simulations, the overall total protein alpha carbon RMSD (Å) decreased with additional Gb3-NBD occupancy, approaching a RMSD of 2.5 Å with two ligands bound. The empty dimer rapidly collapses, via a scissoring motion, where the hairpin ‘roof’ of one chain moves toward the terminal ‘floor’ of the other (Fig. 2d). However, the presence of one or more Gb3-NBD molecules restricts this collapse. Absent any cargo, the binding cavity collapses, and the cavity volume is minimal; but the presence of two Gb3-NBD increases the pocket volume closer to that seen in the crystal structure (Fig. 3a), suggesting that the SapB dimer in the crystal structure contains two or more ligands.

**Figure 3:**
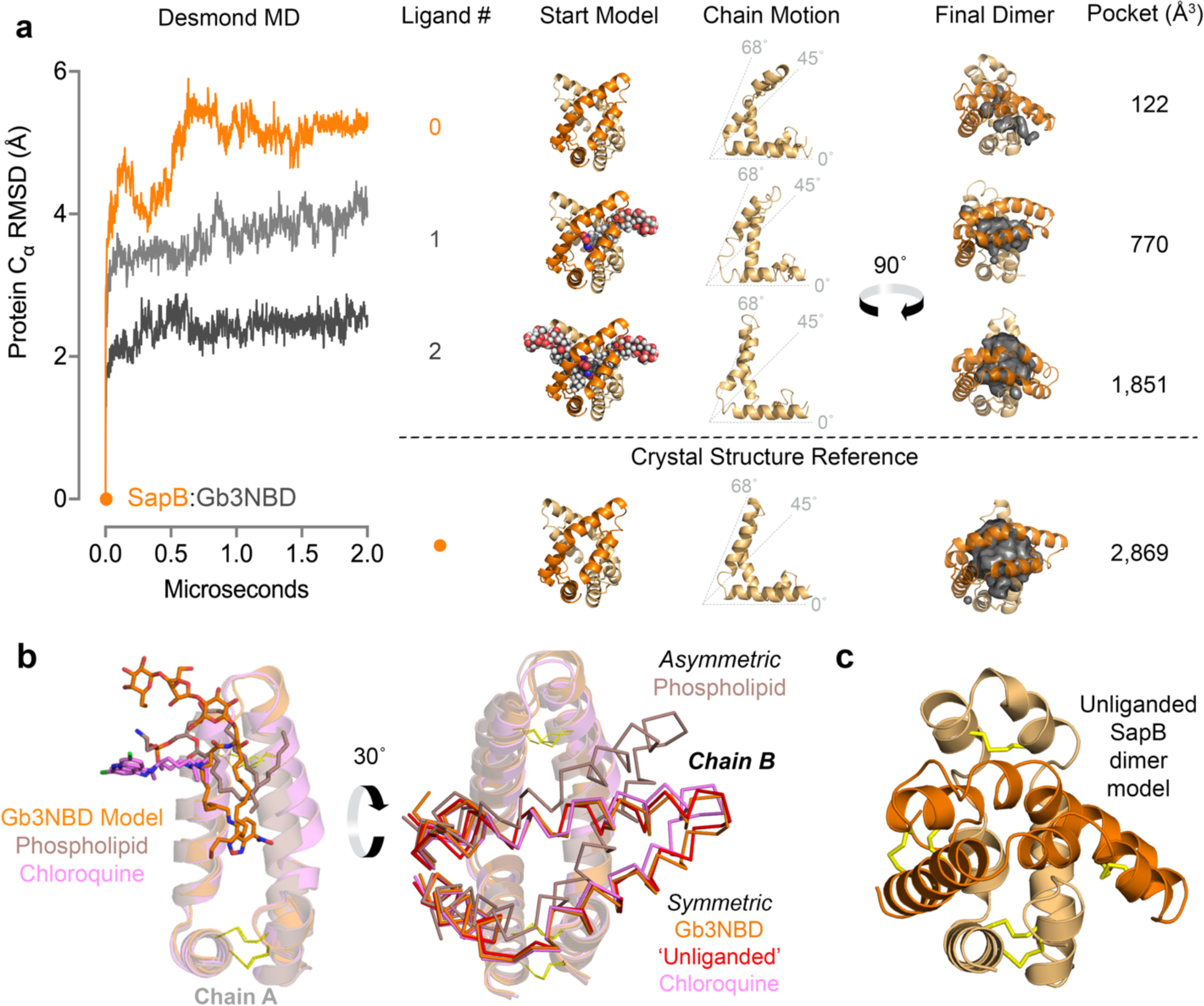
SapB dimer collapses in the absence of ligand. **a** Starting models and resultant movements of empty (orange) and Gb3-NBD-bound SapB dimers (light grey for one Gb3-NBD & dark grey for two Gb3-NBD molecules). The alpha carbon RMSD of the SapB main chains is graphed by microsecond after a MD simulation. Resultant motions in chain B (light-orange) and the final dimer, as a function of ligand presence, are shown for representation. The calculated cavity volumes within the final dimer are indicated to quantify the relative dimer collapse. For reference, the SapB:Gb3-NBD structure (orange dot) is provided under the dotted line. **b** *Left*: An alignment of chain one from the SapB:Gb3-NBD structure (Gb3-NBD modeled ligand, orange stick) to the corresponding chains of other ligand-bound SapB structures (phospholipid, brown; chloroquine, pink) previously published. The Gb3-NBD mimics the location of the resolved ligands of the previous structures. *Right*: A rotated view of the SapB structural alignment, with the second chain rendered (ribbons) to show the symmetry of the dimers. The symmetric ‘unliganded’ structure of SapB (red) was from the same crystal as the asymmetric phospholipid-bound dimer, which had phospholipid present. **c** The unliganded model of a SapB dimer from the MD simulations shown in the orientation of panel b.

As additional controls, we subjected the phospholipid-bound structure of lung surfactant protein (SP-B)^40^, a SapB homolog that binds lipids at acidic pH, to the same simulation (Supplementary Fig. 3a). Absent the phospholipids, a collapse of the hairpin ‘roof’ was also observed, burying the interior hydrophobic residues; the replacement of the phospholipids lowered the RMSD to 2.6 Å, comparable to the SapB model containing two bound Gb3-NBD molecules. Simulations performed on the asymmetric dimer of SapB bound to a single phospholipid^11^ showed a similar collapse in the absence of ligand (Supplementary Fig. 3b). Alignment of the SapB:Gb3-NBD structure to previous SapB structures suggests that the modeled Gb3-NBD, overlaps with the weak experimental difference map density and corresponds to the ligand positions in the homologous structures (Fig. 2c,3b). Moreover, because the chains of the SapB:Gb3-NBD dimer align well with SapB dimers containing ligands, we propose that our structure represents an open and multi-ligand-bound SapB (Fig. 3b). In particular, the open SapB:Gb3-NBD dimer is similar to the dimer of SapB bound to chloroquine^41^ and the ‘unliganded’^11^ crystallographic dimer of SapB in the presence of phospholipid, suggesting that those structures may also contain additional disordered ligands (Fig. 3b). Using SP-B as a reference, which has three resolved phospholipids in the hydrophobic pocket, these SapB structures demonstrate pocket volumes capable of possessing two or more ligands simultaneously (Supplementary Fig. 3c). Overall, the structures from these simulations indicate that absent ligand, SapB has a propensity to bury its hydrophobic pocket, supporting the ligand-bound states of the open SapB dimers in the SapB:Gb3-NBD crystals (Fig. 3c).

### SapB directly interacts with GLA in a ligand-mediated manner *in vitro*

After confirming our loading of SapB with Gb3-NBD, we used chemical cross-linking to capture the transient complex between SapB and GLA with the 11.4 Å cross-linker bis(sulfosuccinimidyl)suberate (BS^3^)^42^, which reacts preferentially with primary amino groups^43^. Using this reagent, we captured a water-soluble ternary complex between a Gb3-NBD-bound SapB and GLA^D170A^. To test the requirement for ligand, we titrated Gb3-NBD against a fixed amount of SapB, GLA, and BS^3^ cross-linker at pH 6.5. Cross-linking produced new bands indicative of a complex between SapB and GLA (Fig. 4a, lanes 10-12). As expected, homodimers of SapB and GLA appeared in the cross-linked samples (lanes 1-8), demonstrating the effective cross-linking of the individual proteins. The intensity of the SapB:GLA complex band captured depends on the presence of Gb3-NBD ligand (Fig. 4a, Supplementary Fig. 4a) and upon the concentration of cross-linker (Fig. 4b).

**Figure 4:**
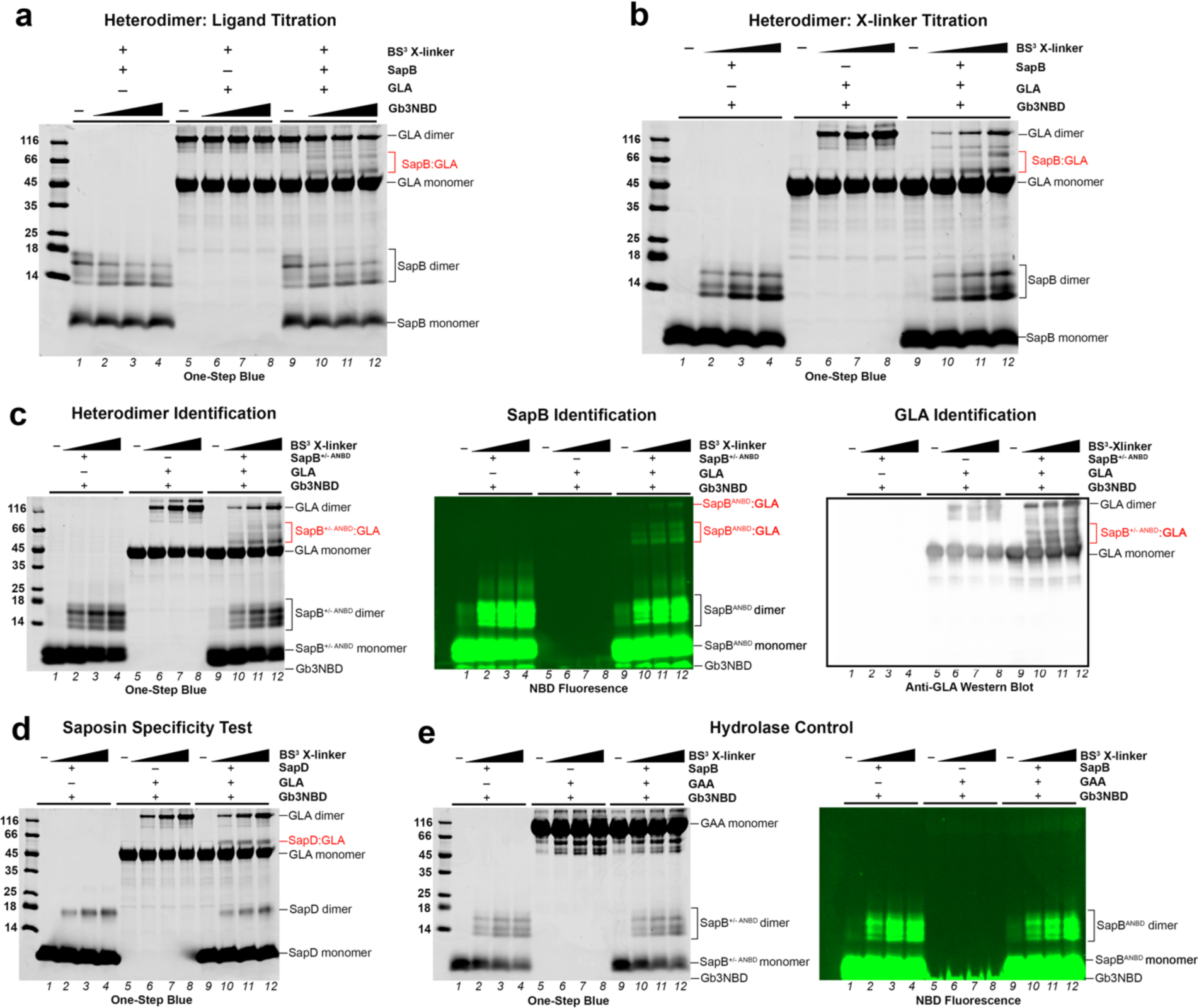
*In vitro* cross-linking of SapB and GLA in the presence of Gb3. **a** One-Step Blue stained SDS-PAGE gel of 10 μM GLA and 50 μM SapB samples after incubation with 1 mM BS^3^ X-linker in the presence of 0, 50, 100, and 200 μM Gb3-NBD. **b** SDS-PAGE gel of 10 μM GLA and 100 μM SapB samples after incubation with increasing concentrations of BS^3^ X-linker (0, 0.2, 0.5, 1 mM) in the presence of 100 μM Gb3-NBD. **c** *Left*: Same as panel b but with 20% reporter SapB^ANBD^ premixed with SapB prior to the cross-linking reaction. *Middle*: The NBD fluorescence of the SDS-PAGE gel prior to One-Step Blue staining to detect the NBD-containing bands using an XcitaBlue conversion screen and GelGreen detection filter. *Right*: An anti-GLA chemiluminescent western blot of duplicate SDS-PAGE samples. **d** Same as panel b but with SapD instead of SapB. **e** *Left*: SDS-PAGE of 10 μM GAA and 100 μM SapB samples (with 20% reporter SapB^ANBD^) samples after incubation with increasing concentrations of BS^3^ X-linker (0, 0.2, 0.5, 1 mM) in the presence of 100 μM Gb3-NBD. *Right*: the NBD fluorescence of the SDS-PAGE prior to One-Step Blue staining.

To verify the identity of the new bands as a SapB:GLA complex, we used an antibody to detect GLA and a covalent, fluorescent reporter of SapB. We expressed and purified a SapB^N21C^ variant (with a reactive thiol replacing Asn21, which is normally N-linked glycosylated) and then labeled it with IANBD amide to generate the fluorescent reporter SapB^ANBD^. This reporter had ∼80% of the activity of wild-type SapB at presenting Gb3-NBD to GLA and measuring galactose production (Supplementary Fig. 1e). In a replicate of the cross-linking assay, the SapB:GLA complex band reacted with an anti-GLA antibody and also fluoresced due to the NBD attached to SapB (Fig. 4c). The Gb3-NBD ligand was undetectable in protein staining but strongly fluoresced at the loading dye front. The heterodimer bands contain SapB^ANBD^ because they both stain as protein and fluoresce (indicating ANBD) (Fig. 4c, Supplementary Fig. 4b).

We first tested the specificity of the SapB:GLA interaction by replacing the saposin and the hydrolase in the cross-linker titration assays. Using SapD in place of SapB led to a SapD:GLA complex band that did not depend on the presence of Gb3-NBD (Fig. 4d, Supplementary Fig. 4c). Second, we replaced the GLA enzyme with acid α-glucosidase (GAA), which does not require SapB as an activator. No complex formed as no SapB migrated larger than the GAA band (Fig. 4e). Third, we lowered the concentration of the presentation molecule and GLA tenfold while maintaining a molar excess Gb3-NBD lipid to test for cross-linking at enzyme-assay concentrations; this cross-linking assay again resulted in discrete SapB:GLA complex bands (Supplementary Fig. 4d). We quantified the solvent accessible BS^3^-reactive residues^43^ in the proteins for reference (Supplementary Fig. 4e). Our results indicate that the SapB:GLA complex depends on the two proteins, the concentration of ligand, and the amount of cross-linker used. Importantly, these results support the hypothesis that there is a transient, direct, and ligand-mediated interaction between SapB and GLA *in vitro*.

### The structure of a SapB:GLA complex

To understand the molecular basis of the SapB interaction with GLA, we grew crystals of SapB:GLA complexes in the presence of Gb3-NBD lipid or decyldimethylamine-*N*-oxide (DDAO) detergent. While crystals of SapB:GLA complexes did not readily form in the presence of the lipid, we did isolate diffraction-quality crystals of a SapB:GLA complex in the presence of DDAO detergent. We determined the structure of SapB bound to GLA at 3.53 Å resolution (Table 2).

**Table 1.** Data collection and refinement statistics for SapB: Gb3-NBD crystals.

|  |  |
| --- | --- |
| **Data Collection** |  |
| Space group | $P 6_4$ |
| Cell dimensions |  |
| $a, b, c$ (Å) | 66.91, 66.91, 87.18 |
| $\alpha, \beta, \gamma$ (°) | 90, 90, 120 |
| Resolution (Å) | 34.83-2.68 (2.78-2.68) |
| $R_{\text{merge}}$ | 0.0432 (0.368) |
| $I/\sigma_I$ | 27.1 (1.51) |
| Completeness (%) | 99.7 (100) |
| Redundancy | 9.7 (10) |
| **Refinement** |  |
| Resolution (Å) | 33.45-2.68 |
| No. unique reflections | 6,241 (609) |
| $R_{\text{work}} / R_{\text{free}}$ | 0.246/0.287 |
| No. atoms |  |
| Protein | 1,157 |
| Other<sup>a</sup> | 10 |
| $B$-factors (Å<sup>2</sup>) | |
| Protein | 94 |
| Water | 84 |
| Ion | 102 |
| R.m.s. deviations |  |
| Bond lengths (Å) | 0.008 |
| Bond angles (°) | 0.97 |
| Ramachandran outliers (%) | 0 |
| Rotamer outliers (%) | 1.6 |
| Clashscore | 2.2 |
| Values in parentheses are for the highest-resolution shell |  |
| <sup>a</sup>Includes water and ions |  |

**Table 2.**
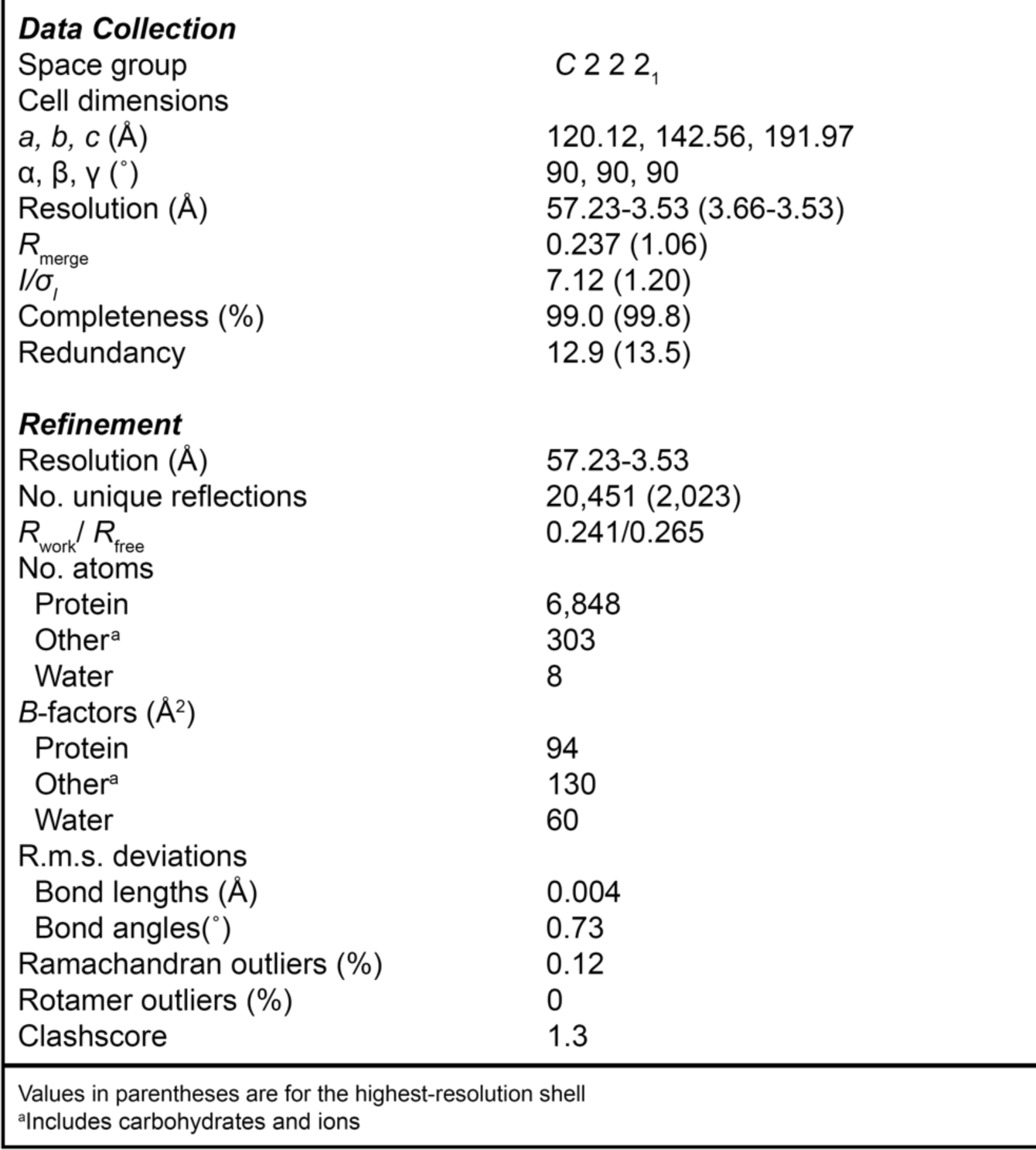
Data collection and refinement statistics for SapB: GLA complex crystals.

|  |  |
| --- | --- |
| Space group | C 2 2 2<sub>1</sub> |
| Cell dimensions |  |
| *a*, *b*, *c* (Å) | 120.12, 142.56, 191.97 |
| $\alpha$, $\beta$, $\gamma$ (°) | 90, 90, 90 |
| Resolution (Å) | 57.23-3.53 (3.66-3.53) |
| $R_{\text{merge}}$ | 0.237 (1.06) |
| $I/\sigma_I$ | 7.12 (1.20) |
| Completeness (%) | 99.0 (99.8) |
| Redundancy | 12.9 (13.5) |
| **Refinement** |  |
| Resolution (Å) | 57.23-3.53 |
| No. unique reflections | 20,451 (2,023) |
| $R_{\text{work}}/R_{\text{free}}$ | 0.241/0.265 |
| No. atoms |  |
| Protein | 6,848 |
| Other<sup>a</sup> | 303 |
| Water | 8 |
| *B*-factors (Å<sup>2</sup>) |  |
| Protein | 94 |
| Other<sup>a</sup> | 130 |
| Water | 60 |
| R.m.s. deviations |  |
| Bond lengths (Å) | 0.004 |
| Bond angles(°) | 0.73 |
| Ramachandran outliers (%) | 0.12 |
| Rotamer outliers (%) | 0 |
| Clashscore | 1.3 |

Values in parentheses are for the highest-resolution shell
<sup>a</sup>Includes carbohydrates and ions

The structure of SapB bound to GLA contains a dimer of GLA and monomer of SapB in the asymmetric unit (Fig. 5a). However, the SapB monomer lies on a crystallographic two-fold axis, generating a symmetry-related SapB pair of monomers resting in the concave architecture of a pair of GLA dimers (Fig. 5b). The SapB pair is proximal to all four active sites of GLA, and the two SapB monomers do not contact each other (Fig. 5b).

**Figure 5:**
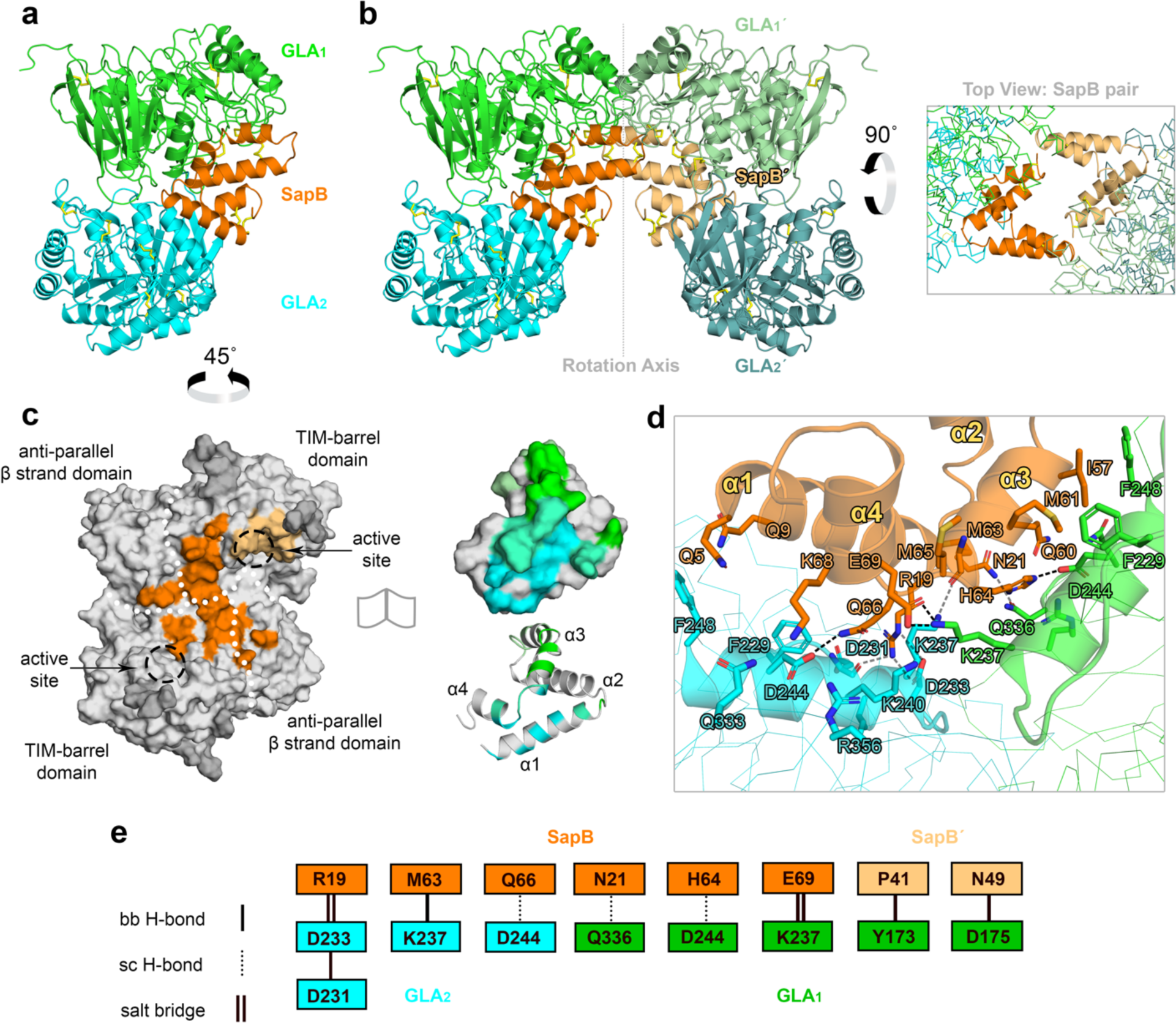
Structure of the SapB:GLA binary complex in the presence of DDAO detergent. **a** Asymmetric unit of the crystallographic complex, consisting of a GLA dimer (green & cyan) bound to a SapB monomer (orange) **b** *Left*: The complex sits near a twofold screw axis that generates a symmetry mate SapB monomer (SapB’, tan) and GLA dimer (GLA_1_’, pale green & GLA_2_’, teal). *Right*: The top view of the SapB pair shows how the SapB chains form crystallographic bridges between the pair of GLA dimers, while the SapB chains do not appear to contact one another. **c** Surface of the GLA dimer (grey) in the asymmetric unit with the SapB interaction sites painted onto it (≤ 4 Å distant residues). The Sap monomer (orange) sits across the dimer interface between the two active sites of both GLA chains and contacts both major domains of the protein (white dot borders). The SapB’ (tan) makes a minor contact near the active site of one chain. Glycans are colored dark grey. GLA-interacting residues (green & cyan) of SapB are conversely shown in both surface and cartoon representations for clarity. The green-cyan blend indicates those SapB residues that touch both GLA monomers. **d** The observed interacting residues (sticks) between several charged, polar, or nonpolar residues of SapB and GLA at the dimer interface of GLA. **e** Schema of the backbone (bb) and side chain (sc) hydrogen bonds and salt bridge interactions observed between SapB (orange), SapB’ (tan), and a GLA dimer (green & cyan).

We painted the SapB footprint onto the GLA dimer to map the complex (Fig. 5c). One SapB monomer rests asymmetrically across the GLA homodimer, contacting each of the two domains of a GLA monomer. The other SapB monomer interacts with one active site of the same GLA dimer. In the context of one GLA dimer and a crystallographic SapB pair, these interactions bury approximately 1,066 Å^2^ (calculated with ePISA^44^), of which 65% is contributed by the one SapB monomer on the GLA dimer interface and 35% is contributed by the other SapB monomer touching the GLA active site. The interacting residues between SapB and GLA at the dimer interface of GLA show charge complementarity, particularly involving the region surrounding the loop between the α3 and α4 helices of SapB (Fig. 5d).

The complex interface contains backbone and side-chain hydrogen bonds and salt bridges, involving Asp231, Asp233, Lys237, Lys240, and Asp244 on GLA and Arg19, His64, Gln66, and Glu69 on SapB (Fig. 5e). The protein complex further reveals van der Waals interactions between Tyr54, Ile57, and Met61 on SapB and Phe229 and Phe248 on GLA. In contrast, the hairpin of SapB, which lies near the active site disulfide bond (Cys142-Cys172) of GLA, interacts largely via van der Waals interactions, with one hydrogen bond between Pro41 on SapB and Tyr173 on GLA.

The structure of the SapB:GLA complex reveals a potential mechanism of glycosphingolipid catabolism and provides a rationale for how some more conservative substitutions in surface-exposed residues on GLA can lead to Fabry disease (Supplementary Fig. 5a). The orientation of the SapB monomers suggests substrate delivery to the active sites of GLA which are near the hairpin of SapB. Thus, we manually modeled the fluorescent substrate Gb3-NBD into the GLA active site, using the ligand conformation of a 1.9 Å crystal structure of GLA bound to melibiose^39^. Not only can each SapB monomer accommodate at least one Gb3-NBD (C12:0 fatty acyl chain) ligand without clash, but there is extensive space for holding at least two native Gb3 (C24:0 fatty acyl chain) molecules within the hydrolase-bound pair (Supplementary Fig. 5b). The complex of SapB:GLA reveals a novel adaptability of the SapB dimer to dissociate into monomers in the presence of lipophilic ligand and a cognate hydrolase.

We combined our structures of SapB:Gb3-NBD and SapB:GLA to generate a model for SapB presenting Gb3 cargo to GLA for hydrolysis. We aligned the SapB:Gb3-NBD structure to SapB in the GLA complex (Supplementary Fig. 5c). This alignment demonstrated that both a 14° narrowing of the SapB hinge angle and a 19 Å separation of the SapB dimer were necessary to form the SapB pair in the SapB:GLA crystal. Accordingly, we infer that SapB dimers separated to form the crystallographic lattice with GLA dimers and the DDAO detergent.

Finally, given that SapD interacts with GLA in cross-linking experiments, we interrogated the specificity of the SapB:GLA complex by the shape complementarity of SapA, C, and D with GLA. Superimpositions of representative structures of SapA–D suggest that only the more compact conformations are compatible (Supplementary Fig. 5d). In contrast, the fully open conformations of saposins result in major clashes with the β strand domain of GLA. These analyses indicate that only more compact saposin shapes are compatible with binding within the concavity of the GLA dimer, suggesting a mechanism for hydrolase selectivity of presentation molecules.

To test the hypothesis that SapB dimers are compatible with presenting to GLA in solution, we performed MD simulations on the SapB:GLA complex to examine the energetics of the association. In each simulation, the GLA dimer showed no appreciable motion, but the SapB changed conformation substantially. Starting with the coordinates of the asymmetric unit containing an open SapB monomer bound at the dimer interface of GLA, we first monitored the translations in the protein chains over 50 ns. The SapB monomer collapsed as expected, resulting in a final RMSD of 3.4 Å for the complex (Supplementary Fig. 6). Second, we ran simulations after adding a single Gb3-NBD molecule in the SapB hydrophobic pocket, having the terminal galactose positioned in the GLA active site. The simulations also showed the SapB collapse, projecting the Gb3-NBD molecule into a hydrophobic channel connecting the SapB interior and GLA active site, with a final RMSD of 2.9 Å (Supplementary Fig. 6). Finally, we simulated the 2:2:2 complex of SapB:GLA:Gb3-NBD. MD simulations on this heterocomplex were the most stable, showing a final 2.6 Å RMSD (Supplementary Fig. 6). These simulations suggest that a compact SapB dimer is more energetically favorable to present cargo to GLA for catabolism.

### Comparison of saposin:hydrolase complexes

We compared the binary complex of SapB:GLA to that of SapA:GALC^15^ and to two lysosomal enzymes with internal saposin domains, acid sphingomyelinase^45^ and acyloxyacyl hydrolase^46^. Aligning the saposins by the floor subdomain of helices α1 and α4 reveals the hydrolase interactions with the α3 helix of the saposins (Fig. 6a). We painted the hydrolase footprints onto the saposin monomers in the structures to examine the differences beyond the α3 helix and found that the enzymes each contact different surfaces on their cognate saposins (Fig. 6b). We also calculated the electrostatic surface potentials of saposins at lysosomal pH and highlighted the saposin residues that would interact with GLA based on the interface formed with SapB in the crystal structure (Fig. 6c). The SapB:GLA specificity is driven by shape and charge complementarity. Sequence alignments of saposins (Fig. 6d) show that residues that bind cognate hydrolases are not conserved, another indication of the specificity of hydrolases for different saposins^15^. Altogether, the quaternary structure, monomer conformation, shape, and charge complementarity of the saposins drive specificity for a partner hydrolase.

**Figure 6:**
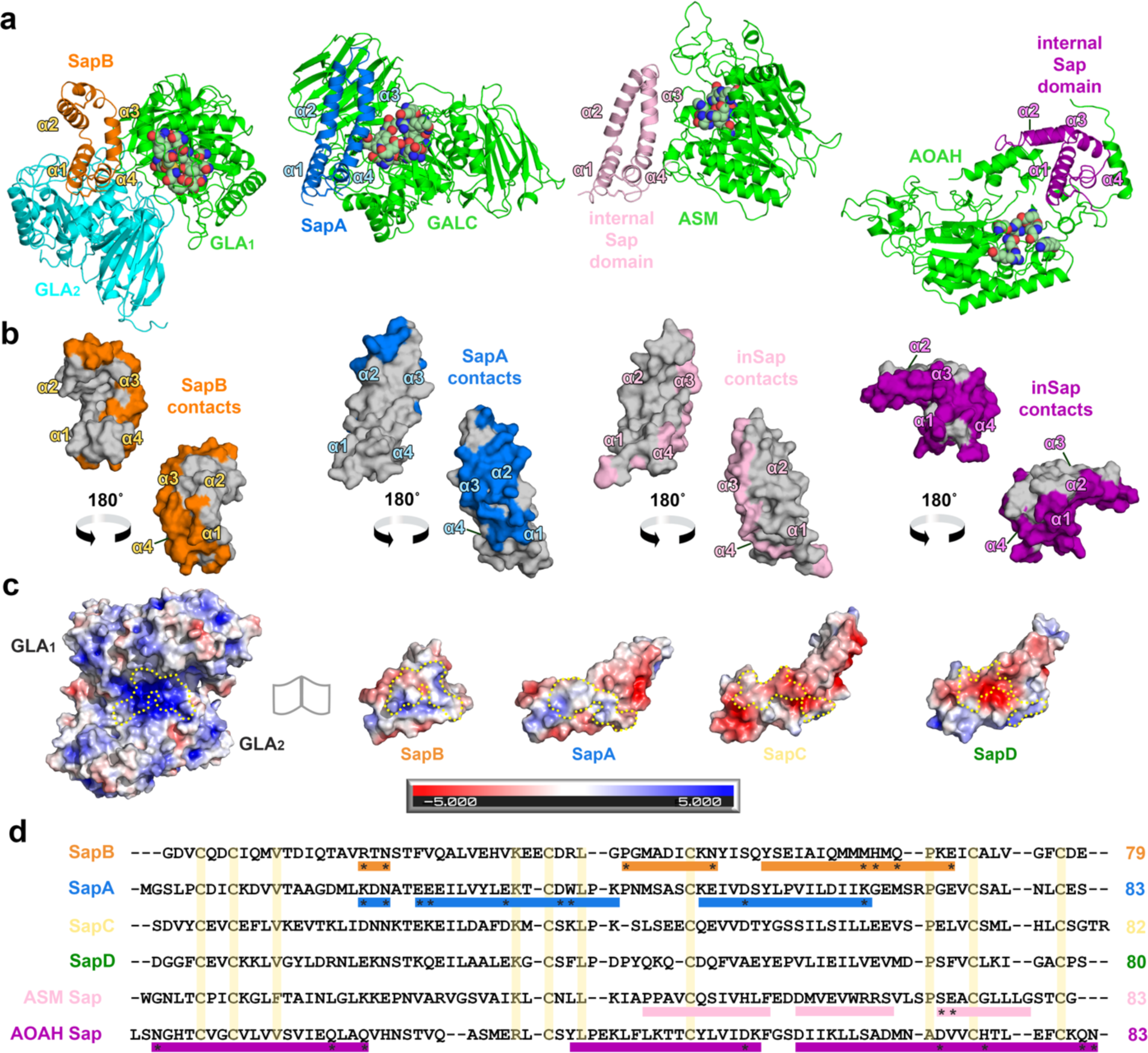
Arrangement and specificity of lysosomal saposin-hydrolase interactions. **a** Comparison of the SapB:GLA, SapA:Galactosylceramidase (GALC), acid sphingomyelinase (ASM), and acyloxyacyl hydrolase (AOAH) crystal structures. The hydrolase chains are shown in green and cyan. The lysosomal saposin (SapB-orange; SapA-blue) or internal saposin domains (ASM-pink; AOAH-purple) are aligned. The most proximal active site is rendered with atomic-colored spheres. **b** Surface of the aligned saposin monomers with the hydrolase-interacting (≤ 4 Å distance residues) residues painted according to the saposin color in panel a. The helices are labeled for orientation. **c** Electrostatic surface potential of GLA and the aligned saposin monomers calculated at pH 5. Red and blue surfaces correspond to negative and positive electrostatic potential scaled from −5 *k_b_T* to 5 *k_b_T*. The major SapB-interacting grove (yellow dotted line) of GLA exhibits a net positive charge whereas the saposins show distinct shapes and charge distributions. Residues on the other saposins which align with the GLA-interacting residues of SapB are similarly outlined. **d** Multiple sequence alignment of the lysosomal saposins and internal saposin domains of hydrolases. The hydrolase-interacting regions are shown (colored rectangles) with residues forming key electrostatic interactions indicated (asterisks). The conserved residues (> 80 % identity) including the disulfide-bond forming cysteines are highlighted (yellow).

## DISCUSSION

Here we describe the transient ternary interaction among the lysosomal hydrolase GLA, activator protein SapB, and the ligand Gb3-NBD, a fluorescent analog of the natural substrate Gb3 that accumulates in Fabry disease patients. These results address the mechanistic basis for processing the Gb3 substrate and for the failure of processing that leads to Fabry disease. For example, prior to our results, it was unclear how conservative substitutions in surface residues like D244N and D175N could lead to the complete loss of enzymatic activity that characterizes Fabry disease. Structural analysis of the SapB:GLA complex suggests that those substitutions affect the transient interaction between GLA and SapB that is required for the hydrolysis of Gb3.

We used chemical cross-linking to trap transient complexes between SapB and GLA. These cross-linking experiments allowed us to identify intermolecular interactions that were too short-lived for other biochemical assays. However, gel-filtration chromatography allowed the tracking of longer-lived lipoprotein complexes containing the fluorescent Gb3-NBD, where one wavelength (280 nm) tracked protein and another (460 nm) tracked fluorescent sphingolipid. The fluorescent Gb3-NBD was critical for detecting ligand in SapB crystals in the absence of ordered electron density. These results substantiate the rationale for employing NBD-reporter lipids in other binding studies.

Our biochemical assays show that SapB binds Gb3-NBD and activates GLA. Surprisingly, we also detected some SapD association with GLA. The SapD activation of GLA was 7% as efficient as SapB, suggesting that SapD may interact with GLA under some circumstances. The capture of SapD:GLA heterocomplexes in the cross-linking assays suggests the possibility of inter-saposin competitive inhibition, which phenomenon has been posited to influence lipid substrate availability and the magnitude of hydrolase activation^47,48^. As such, conducting cross-linking assays with the four saposins and their cognate hydrolases may help to define the scope and selectivity of these transient interactions.

Our crystal structures and MD simulations reveal surprising quaternary interactions in SapB. We show that the open conformation of SapB seen in previous several crystal structures is likely to have ligand bound. Despite the ambiguous electron density in the hydrophobic cavity of the SapB:Gb3-NBD crystals, we show by optical imaging of the crystals, fluorescence spectroscopy, mass spectrometry, and molecular dynamics that the SapB structure binds Gb3-NBD cargoes. In the Gb3-NBD and Cer-NBD fluorescent binding assays, the addition of iodide failed to quench the NBD fluorescence suggesting that the NBD moiety is solvent inaccessible in the hydrophobic pocket of SapB. Computational modeling indicates that in the absence of ligand, the SapB dimer collapses into a compact globular shape that buries its exposed hydrophobic residues. The collapse of SapB in the absence of ligand was previously suggested by MD simulations on SapB^49^. We suggest that previous crystal structures of SapB in open conformations likely contain bound but disordered ligands. Our simulations indicate that an open SapB dimer binds at least two Gb3-NBD molecules. Furthermore, these results are consistent with reports describing *in vitro* experiments showing SapB dimers binding two phospholipids^50^ and *in silico* experiments showing SapB dimers binding two small molecules^51^.

The SapB:GLA complex structure shows that the SapB dimer adapts to different ligands, as expected for a carrier that presents hydrophobic ligands to hydrolytic enzymes. The conformation of SapB in the SapB:GLA complex immediately suggests a mechanism for how SapB presents Gb3 to GLA for cleavage of the terminal galactose from Gb3. The SapB:GLA structure also explains the concave shape of the human GLA dimer. The concave hydrolase complements the convex shape of SapB, and SapB binds directly on the molecular dyad of GLA. The concave shape of the GLA dimer allows SapB to present for cleavage a wide variety of ligands with terminal α-galactose saccharides. Beyond binding to Gb3, SapB forms soluble complexes with branched detergents and lipids, including lactosylceramide and galactosylceramide^52,53^.

The transient complexes between mature lysosomal saposins and their cognate hydrolases promote glycosphingolipid catabolism, but the transient complexes are difficult to characterize structurally. Our structure of a SapB:GLA complex now joins the structure of a SapA:GALC complex^15^ to lead to general principles for saposin:hydrolase interactions (Fig. 6). In both complexes, the saposin α3 helix appears near the hydrolase active site with the solvent-exposed N-linked glycosylation sites remaining facing away from the hydrolase. This geometry indicates that (a) *E. coli* expressed saposins can form relevant complexes with their cognate hydrolases and (b) substitutions at the N-linked glycan site, like our SapB^N21C^ and the previously reported SapC^N22C^,^54^ can yield reporters for exploring saposin:hydrolase associations. Combined with the structures of other hydrolases containing internal saposin domains (Fig. 6), we now have a more comprehensive basis for understanding ligand and hydrolase specificity in the saposin family.

Our MD simulations and the SapB:Gb3-NBD structure, as well as previous reports on SapB, suggest that the SapB dimer binds lipids and presents cargoes for cleavage^8,11,50^. While most evidence supports a dimer as the functional quaternary structure of SapB, we cannot exclude the possibility that transient, functional SapB monomers, as captured in the crystal of the SapB:GLA complex, may modulate GLA activity. Because SapA, SapC, and SapD can change quaternary structures^5–8,55,56^, it remains possible that SapB does also.

Crystal structures capture snapshots of dynamic molecular processes^57^. For instance, the bent ‘leg’ conformations observed in the first crystal structure of the ectodomain of integrin αVβ3,^58^ inspired further research concerning their physiological relevance^59^. For example, activation of αVβ3 with manganese ions was found to induce αVβ3 molecules to adopt extended leg conformations in negative stain electron microscopy experiments as well as increase ligand affinity^60^. However, another study of manganese-bound, native-sequence αVβ3 ectodomain:fibronectin soluble complexes presented the bent conformation as the major species of particles in electron micrographs^61^. Ligand-competent, bent conformations were next observed in a crystal structure of the complete αVβ3 ectodomain and an α/β transmembrane fragment^62^ and, more recently, in a cryo-EM structure of a homologous, full-length integrin αIIbβ3 in the presence of native lipids^63^. These reports illustrate how an unusual conformation of a protein in a crystal can combine with modeling and other supporting information to lead to a broader understanding of its function.

The crystal structure of the SapB:GLA complex presented here, combined with the recently determined SapA:GALC complex,^15^ together provide insight into the underlying principles of interactions between mature lysosomal saposins and their cognate hydrolases, including their ligand dependence and quaternary structures. More research is required to fully understand the rules for saposin-hydrolase molecular interactions.

Overall, our findings are supported by a range of orthogonal experiments, including fluorescence spectroscopy, chromatography, enzyme kinetics, mass spectrometry, X-ray crystallography, chemical cross-linking, and MD simulations. Together, these experimental approaches combine to provide a robust understanding of the role of SapB adaptability in its broad range of cellular functions.

## METHODS

### Materials

N-Dodecanoyl-NBD-ceramide trihexoside and N-dodecanoyl-NBD-D-*erythro*-dihydrosphingosine lipid analogs were purchased from Matreya (Pleasant Gap, PA, USA). 2,2-didecylpropane-1,3-bis-β-D-maltopyranoside (NG310) detergent was purchased from Anatrace (Maumee, OH, USA). Egg white lysozyme (BP535-1), BS^3^ (bis(sulfosuccinimidyl)suberate crosslinker (A39266), IANBD amide (*N*,*N*’-dimethyl-*N*-(iodoacetyl)-*N*’-(7-nitrobenz-2-oxa-1,3-diazol-4-yl)ethylenediamine) (D2004), and the Amplex Red galactose assay (A22179) were purchased from ThermoFisher Scientific (Waltham, MA, USA). The synthetic α-galactosidase substrate *p*-nitrophenyl-α-D-galactopyranoside (N0877) was purchased from Sigma-Aldrich (St. Louis, MO, USA). Human acid α-glucosidase was a gift from Genzyme (Cambridge, MA, USA). Sodium malonate was purchased from Hampton Research (Aliso Viejo, CA, USA).

### Protein Expression and Purification

*E. coli* expressed saposins: We purchased from Genscript (Piscataway, NJ, USA) the codon-optimized SapB and SapD (UniProt P07602) nucleotide sequences inserted between the *Nde*I and *Hin*dIII sites of the expression vector pET-21a. Genscript performed site-directed mutagenesis on the SapB plasmid to make the mutant SapB^N21C^, providing a single free cysteine mutant at the glycosylation site. The saposin sequences had enterokinase-cleavable N-terminal His8-DDDDK-GAS tags to expediate purification. The plasmids were transformed into *E. coli* Shuffle T7 Express cells (C3029J, New England Biolabs) to aid disulfide bond formation. The expression and purification protocols were adapted from literature precedents^41,64^. In detail, transformed cells were grown at 30 °C from a starter culture in Terrific Broth with carbenicillin and then scaled up to a 1 L baffle flask with ampicillin. When the OD600 reached 0.4-0.8, expression was induced with the addition of isopropyl β-D-1-thiogalactopyranoside to a final concentration of 1 mM. The temperature was lowered from 30 °C to 20 °C and the culture was incubated overnight. Cells were then harvested by centrifugation at 4,000 *g* for 20 min at 4 °C. Cells were then resuspended in His-Trap binding buffer (50 mM Tris pH 7.4, 150 mM NaCl) and pulse sonicated for up to 12 min over ice. Lysates were spun at 4,000 *g* for 45 min at 4 °C and supernatants were recovered. The samples were heat shocked in a water bath for one hour at 95 °C and then re-centrifuged to remove precipitated protein. The heat-shocked supernatants were filtered through a 0.45 μm polyethersulfone (PES) membrane prior to the addition of ProteIndex Ni-Penta agarose (Marvelgent Biosciences) for batch binding at 4 °C for one hour. The resin was collected in a gravity column and washed with 5 column volumes of binding buffer with 4 M urea to favor release of endogenous co-purifying lipids in the samples. After washing with binding buffer to remove the urea, a one-step elution from the Ni^2+^ column was carried out with His-trap elution buffer (50 mM Tris pH 7.4, 150 mM NaCl, 250 mM imidazole). The eluted sample was dialyzed into enterokinase cleavage buffer (20 mM Tris pH 7.4, 50 mM NaCl, 2 mM CaCl2) over a HiPrep 26/10 desalting column (Cytiva) according to the manufacturer’s instructions. His-tagged Enterokinase (Z03004, Genscript) was added to the dialyzed, pooled samples and cleavage was conducted for 24 hours at room temperature and then 24 hours at 4 °C. The resulting solution was then separated in an ÄKTA pure 25 FPLC system (Cytiva) with a 5 mL ProteIndex Ni-Penta column at flow rate of 1 mL/min, and the flow through containing the untagged saposin was collected. The saposin sample was concentrated on a 3-5 kDa MWCO PES membrane Vivaspin 20 Centrifugal Concentrator (Sartoris) and loaded onto a Superdex 200 10/300 GL (Cytiva) size-exclusion chromatography column equilibrated in storage buffer (50 mM Tris, 150 mM NaCl, pH 7.8) at a flow rate of 0.5 mL/min and injection volume of 300 μL in a 500 μL loop. The major peak fractions containing pure saposin were collected and concentrated to ∼5-15 mg/mL for storage at 4°C, with 0.01% azide. Before all assays, the saposin protein was spun at 16,000 *g* for 10 min and buffer exchanged over a Zeba spin desalting column (ThermoFisher) with a 7 kDa MWCO into the appropriate buffer.

Sf9 expressed GLA: We purchased from Genscript the codon-optimized wild-type and D170A mutant human *GLA* (UniProt P06280) nucleotide sequences inserted between the *Csp*I and *Hin*dIII sites of the expression vector pFastBac1. The GLA sequences had a C-terminal 2x-FLAG tag to expedite purification from the supernatant. The plasmid was transformed into the DH10Bac *E. coli* strain (ThermoFisher), blue-white colony screen was performed, and viral infection and amplification in Sf9 cells (Expression Systems) was conducted using Bac-to-Bac Baculovirus Expression System (Invitrogen) protocols. In detail, 1 μg of purified bacmid plasmid DNA was mixed with 8 μL of Cellfectin II (ThermoFisher) at room temperature to transfect 0.9×10^6^ Sf9 cells per well in a 6-well tissue culture plate in a semi-humid environment. All subsequent incubation steps were conducted at 27 °C. After 4-5 hours, the transfection media was exchanged for ESF-921 media (Expression Systems, Davis, CA, USA), and the culture was incubated for 96 hours, after which the P0 viral supernatant was harvested and filtered over a 0.2 μm PES membrane syringe filter. Viral amplification in log-phase suspension Sf9 cultures was performed using a multiplicity of infection (MOI) of 0.1 with harvesting and filtering of the viral supernatant after 72 hours. Once the P2 virus was obtained, Sf9 cultures were scaled up to 400 mL and infected in log-phase with an MOI of 1-3. After incubation for 72 hours, the cells were spun down at 4000 *g* for 30 min at 4 °C and GLA protein was purified by batch binding with 2.5 mL of Anti-DYKDDDDK G1 Affinity Resin (L00432, Genscript) in the harvested and 0.8 μm filtered supernatant. Binding was carried out at 4 °C for at least one hour, and the resin was gravity precipitated and put into a gravity column. The resin was washed with ≥ 20 column volumes of Phosphate Buffer Saline (PBS; ∼10 mM sodium phosphate, 1.8 mM potassium phosphate, 2.7 mM potassium chloride, 137 mM sodium chloride; pH by hydrochloric acid titration) pH 6.5 prior to competitive elution with ∼2.5 column volumes of buffer containing 0.3 mg/mL of DYKDDDK tag peptide (A6002, APExBIO). The eluted protein was concentrated on a 10 kDa MWCO PES membrane Vivaspin 6 Centrifugal Concentrator (Sartoris) to ∼4.7 mg/mL and stored at 4 °C, with 0.01% azide. Diagnostic SEC runs were conducted on a Superdex 200 10/300 GL column equilibrated in PBS pH 6.5 buffer at a flow rate of 0.5 mL/min to quality-check protein lots. Before all assays, the GLA protein was spun at 16,000 *g* for 10 min and buffer exchanged over a Zeba spin desalting column (ThermoFisher) with a 7 kDa MWCO membrane into the appropriate buffer, which also removed leftover FLAG peptide.

### NBD Fluorescence Binding Assays

SapB, SapD, or lysozyme was diluted into 50 mM MES pH 4.6 or 6.5 buffer to a final concentration of ∼0.33 mM in a volume of 140 μL per replicate (n=3) in a black flat-bottom non-binding microplate well. A 16-fold serial 1:1 dilution series was made by carrying over 70 μL of each protein sample and mixing it with 70 μL of the buffer. Subsequently, 2 μL of a ∼80 μM stock of Gb3-NBD or Cer-NBD (in 100% DMSO) was spiked into each protein well, resulting in ∼2.2 μM lipid. The plate was incubated for one hour at room temperature and endpoint fluorescence intensities were read 3 times at 30 °C with a BioTek Synergy H1 plate reader using 460 nm excitation and 540 nm emission wavelengths. On a separate plate, several lipid- and buffer-alone samples were run in parallel. The protein background fluorescence was detected prior to the addition of lipids. After the one-hour timepoint, plates were sealed and left to incubate for 21 days at room temperature and then read again. For the quenching of NBD, sodium iodide was diluted into each well to a final concentration of 27 mM and, after one hour, the plates were read a final time. Binding curves were fit using the nonlinear regression *log (agonist) versus response – variable slope* function in GraphPad Prism 9 Software (San Diego, CA, USA).

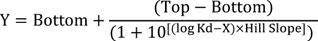

The Kd is the concentration of the activator (X-axis) that gives a response halfway between the Bottom and Top. The plateaus of the binding curve are measured in relative fluorescent units (RFU; Y-axis). The Hill Slope describes the steepness of the sigmoidal curve between the plateaus and is indicative of the binding dependency of the activator to the substrate molecule. Estimating a standard saturation binding model between protein and lipid, we report the apparent dissociation constant, Kdapp, consistent with the experimental setups.

### Size-Exclusion Chromatography Studies

A 10% volume of Gb3-NBD (10 mg/mL; 100% DMSO) was added to a sample of ∼1 mM SapB, SapD, or lysozyme in PBS 6.5, resulting in an approximate 1:1 molar ratio of Gb3-NBD:protein. The samples were incubated at 37 °C for one hour and then 50 μL was injected onto a Superdex 200 10/300 GL column at a flow rate of 0.5 mL/min with an AKTA pure 25 FPLC system with a PBS pH 6.5 mobile phase. The absorbances at 280 nm and 460 nm were monitored with a U9-M detector. Equivalent protein alone samples with DMSO solvent additions were run as controls. A 5% volume of Gb3-NBD stock was added to the buffer and run as the lipid-alone baseline. For a non-protein carrier molecule sample, a 5% volume of Gb3-NBD was added to a 52.5mM lauryl maltose neopentyl glycol (LMNG) detergent sample. To approximate the magnitude of Gb3-NBD as a function of carrier molecule, the *area under the curve* function in GraphPad Prism 9 was used and compared with the theoretical total area of a 50 μL sample of the lipid alone in buffer with the absorbance measured at 460 nm.

### Enzyme Kinetics

GLA activity was determined using the synthetic substrate *para*-nitrophenyl-α-galactose (pNP-α-Gal) in 150 mM citrate-phosphate pH 5 buffer using substrate concentrations 0, 1, 3, 5, 10, 20, 40, and 50 mM (n=3). GLA was added at a final concentration of 0.01 mg/mL to the pNP-α-Gal, and reaction aliquots were taken every 1.5 min for 10.5 min and diluted into 200 mM borate buffer pH 10 in a clear flat bottom plate to stop the reaction. Absorbance measurements at 405 nm were taken to determine the concentration of pNP released from α-Gal, using Beer-Lambert law with the molar absorption coefficient of 18.1 mM^-1^cm^-1^. Km, Vmax, and kcat were calculated by fitting the data to a Michaelis-Menten equation in GraphPad Prism.

### Enzyme-Activator Assay

Enzyme-activator samples were prepared in 50 μL reactions by mixing 0.1 mg of BSA, ∼11 μg Gb3-NBD, 2 μg of wild-type GLA, and either 0.04, 0.4, or 4 μg of SapB in PBS pH 4.4 in a black flat-bottom non-binding microplate well (n=3). The reactions were incubated at 37°C for 2.5 hours, after which the free galactose was detected with the Amplex Red Galactose/Galactose Oxidase Kit (Invitrogen) according to the manufacturer’s instructions. Briefly, the enzyme-activator reaction was mixed 1:1 with a 2X working solution of 0.1 mM Amplex Red reagent containing 0.2 U/mL horseradish peroxidase and 4 U/mL galactose oxidase in 50 mM Tris-HCL pH 7.2 and 5 mM CaCl2. After a 30 min incubation at 37°C, the fluorescent endpoint intensities were read six times with a BioTek Synergy H1 plate reader using 560 nm excitation and 590 nm emission wavelengths to detect the red fluorescent resorufin which is generated by Amplex Red reacting 1:1 with hydrogen peroxide coming from the oxidation of free galactose. To establish the floor and ceiling of GLA activity, SapB was omitted, or 0.27% sodium taurocholate detergent was substituted in place of SapB, respectively. SapB was replaced by SapD for a specificity control. As a control, the catalytic nucleophile mutant GLA^D170A^ was utilized. Additionally, saposin alone, Gb3-NBD alone, and hydrogen peroxide alone were incubated overnight with the 2X working solution as controls. To calculate the concentration of free galactose in the samples, a standard curve was made in parallel by making a 1:1 serial dilution of galactose alone in buffer ranging from 0–120 μM.

### Crystallization, X-ray Data Collection, and Structure Determination

*SapB:Gb3-NBD crystals*: SapB was concentrated to ∼8.3 mg/mL in storage buffer. A 10% volume of Gb3-NBD (10 mg/mL, 100% DMSO) was added to the SapB, resulting in ∼7.5 mg/mL SapB (∼0.82 mM) and 1 mg/mL Gb3-NBD (0.87 mM), an approximate molar equivalence. The sample was incubated for 2 hours at 37 °C in an Eppendorf Protein LoBind tube. Sitting drop vapor diffusion crystallization trials were performed at the nanoliter scale (500 nL protein and 250 nL crystallization reagent) using an NT8 Drop Setter (Formulatrix) on Greiner CrystalQuick Plus hydrophobic plates. Diffraction quality crystals grew at 20 °C within a fine grid screen after ∼3 months using 3.1-3.3 M malonate pH 4.9-5, of which the best was grown using 3.25 M malonate pH 5. Crystals were harvested and directly flash-cooled by plunging into liquid nitrogen. Diffraction data were collected at the Stanford Synchrotron Radiation Lightsource (SSRL) on beamline 12-2 with a Dectris Pilatus 6M detector. Diffraction data were indexed and integrated with *XIA2*^65^ *DIALS*^66^ within the CCP4i2 (v.7.1)^67^ suite. Molecular replacement was performed by searching for 2 copies of a single SapB polypeptide chain (PDB ID: 4V2O) in the asymmetric unit using *phenix.phaser*^68^ in Phenix (v. 1.19.2)^69^, and refinement was carried out using *phenix.refine*^70^ using translation-libation-screw parameters and non-crystallographic symmetry restraints. Manual rebuilding and map inspection was performed in *Coot* (v. 0.9.5)^71^, and structure figures were generated using PyMOL (v. 2.4.1, Schrödinger, LLC). The crystallographic data collection and refinement statistics for SapB:Gb3-NBD crystals are presented in Table 1. *Structure analysis*: Structural alignments utilized the SapB N- and C-terminal residues (#1-23, 67-78) as a template reference. For NBD-fluorescence emission detection, several dozen crystals were individually looped, twice-washed (∼15 sec in 3.4 M malonate pH 5) and dissolved in PBS pH 6.5, alongside a crystal buffer control using the BioTek Synergy H1 settings above.

*SapB:GLA crystals*: SapB (15 mg/mL in 25 mM Tris pH 7.8, 25 mM NaCl) was diluted to 0.22 mM in acidic buffer (50 mM acetate pH 5, 700 mM NaCl) and spiked with 10.5 mM DDAO (105 mM stock in Mili-Q water) resulting in a ∼1:47 SapB:DDAO molar ratio. The SapB:DDAO sample was incubated for 2 hours at 37 °C in an Eppendorf Protein LoBind tube before mixing 1:1 (v/v) with 0.1 mM GLA^D170A^ (PBS pH 6.5) for one hour at room temperature. Immediately before allocation into the crystallization condition, the complex sample was mixed 1:1 with Mili-Q water. Sitting drop vapor diffusion crystallization trials were performed at the nanoliter scale (500 nL protein and 500 nL crystallization reagent using an NT8 Drop Setter (Formulatrix) on Greiner CrystalQuick Plus hydrophobic plates.

Diffraction quality crystals grew at 20 °C within 2 weeks using 0.1 M Tris pH 8, 1 M ammonium sulfate. Crystals were harvested and directly flash-cooled by plunging into liquid nitrogen after cryoprotection with 10% glycerol, 20% PEG 400, and 10.5 mM DDAO added to the crystallization reagent. Diffraction data were collected at the SSRL on beamline 9-2 with a Dectris Pilatus 6M detector. The data were indexed and integrated with *XIA2 DIALS* within the CCP4i2 suite. Molecular replacement was performed by searching for 2 copies of a single GLA polypeptide chain (PDB ID: 3S5Z; non-protein atoms removed) in the asymmetric unit using *phenix.phaser*, and refinement was carried out using *phenix.refine* using non-crystallographic symmetry restraints. The 2mFo-DFc and mFo-DFc maps confirmed the D170A mutation and SapB presence. The SapB chain was built piecemeal with fragments from previous SapB structures. Manual rebuilding and map inspection was performed in *Coot*, and structure figures were generated using *PyMOL*. The crystallographic data collection and refinement statistics for the SapB:GLA complex are presented in Table 2. *Structural analysis*: Structural alignments utilized the SapB N- and C-terminal residues (#1-23, 67-78) or the α2-hairpin-α3 residues (#24-63) as the reference. The PDB coordinates for SapA (ID: 2DOB and 4DDJ), SapC (ID: 1SN6, 2GTG, and 2QYP), and SapD (ID: 3BQQ and 5U85) were used for inter-saposin comparisons. Additionally, the coordinates for SapA:Galactosylceramidase (ID: 5NXB), acid sphingomyelinase (ID: 5I81), and acyloxyacyl hydrolase (ID: 5W78) were used for comparison of crystallized saposin:hydrolase complexes. The angles of the SapB hinges were calculated using the *DynDom* server. The Adaptive Poisson-Boltzmann Solver (*APBS*) software^72^ and associated *PDB2PQR*^73^ servers were used to calculate the electrostatic surface potentials of the hydrolases and saposins. The sequence alignment of the saposins was conducted using MUSCLE^74^ within SnapGene software (v. 4.3.1.1).

### Liquid Chromatography Mass Spectrometry

Several dozen SapB:Gb3-NBD crystals were individually looped and twice-washed in 3.4 M malonate pH 5 buffer and dissolved in Mili-Q water, yielding a final absorbance of 0.03 at 280nm (∼ 0.82 mg/mL). Samples were diluted to 0.1 mg/mL with 0.1% formic acid, and 10 μl was injected onto a Grace Vydac 214TP C4, 2.1 x 50 mm column (Hichrom, Leicestershire, UK) connected to an Agilent 1100 HPLC system with a binary pump. The mobile phases were (A) 0.1% formic acid and (B) 0.1% formic acid in 99% acetonitrile. The system was equilibrated in 10% B at a flow rate of 250 μL/min and the following gradient was applied: 10% B for 1 min, 10-60% B over 5 min, 60-90% B over 2 min, 90% B for 1min. The flow was infused into a 7 T solariX FTICR mass spectrometer (Bruker, Billerica, MA) equipped with a standard electrospray source. The capillary voltage was 4.5 kV, dry gas flow of 8 L/min and dry gas temperature of 200 °C. MS spectra were acquired over 200 – 3,000 m/z with a 256 Kword transient size and 0.2 sec accumulation time. Data were processed using DataAnalysis Version 5.0 (Bruker).

### Molecular Dynamics

*SapB:Gb3-NBD model building and molecular dynamics*: Zero, one, and two Gb3-NBD molecules were built into the SapB:Gb3-NBD structure using *Coot*. The control protein PDB files for SapB bound to phospholipid (ID: 1N69, chain A-B dimer) and mouse surfactant protein B (SP-B) bound to phospholipid (ID: 6VYN, chain D-F dimer) were extracted with and without ligand. All structures in their PDB formats were loaded serially into the Maestro graphical interface for *Desmond*^75^ molecular dynamics (MD) simulations (Schrödinger v 2020-4). PDB files were preprocessed using the Protein Preparation Wizard to add hydrogens and missing side chains and to assign bond orders and disulfide bonds. Protonation states of the amino acids were determined by *PROPKA*^76^ at the lysosomal pH of 4.5. Waters were deleted and restrained minimization was applied using *OPLS3e*^77^ force field parameters. Before the MD simulation, the system was first built using the *TIP3P*^78^ water model in a minimized orthorhombic box with 0.15 M NaCl and neutralizing Na^2+^ ions. The MD simulation was carried out for 2 microseconds with the system at 300 K and a pressure of 1.01325 bar. The trajectories were analyzed using the built-in interaction diagram module to extract the root means square deviation (RMSD) ‘.dat’ file, and then the trajectories were loaded into *PyMOL* to output the first and last frame PDB files for analysis and figure generation. Cavity pocket analysis was performed with *CASTp*^79^ using a 1.0 Å radius probe. *PyMOL* analyses additionally included the coordinates for SapB bound to chloroquine (ID: 4V20, chain A-B dimer) and SapB bound to phospholipid (ID: 1N69, chain C-C’ dimer).

*Molecular dynamics of the SapB:GLA complex in the crystal structure*: 50 nanosecond MD simulations were performed on the asymmetric unit containing an empty SapB monomer and GLA dimer after preprocessing using the Protein Preparation Wizard as abovementioned. To show the effect of ligand, a single Gb3-NBD molecule was manually built into the SapB monomer using *Coot*, and the terminal galactose was placed in the GLA active site, according to the substrate bound structure of GLA (PDB ID: 3HG3), prior to the simulation. For the ligand-bound dimer, the two-Gb3-NBD modeled SapB:Gb3-NBD structure was aligned to the SapB monomer via the N- and C-terminal residues (#1-23, 67-78). The trajectories were analyzed using the built-in interaction diagram module to extract the RMSD ‘.dat; file and first and last PDB frames as above for analysis in *PyMOL*.

### Fluorescent Labeling of SapB

SapB^N21C^ was concentrated to ∼1.2 mM in PBS pH 6.5 and a 10% volume of pH 7 TCEP (1 M) was added to the SapB, resulting in a ∼1:10 SapB:TCEP molar ratio. After a 20 min incubation at room temperature, the sample was injected onto a Superdex 200 10/300 GL column at a flow rate of 0.5 ml/min with a PBS pH 6.5 mobile phase. Peak fractions containing SapB^N21C^ were pooled and adjusted to ∼0.11 mM as necessary. Next, HEPES pH 7 was added (25 mM final) and semi-soluble IANBD Amide (Invitrogen) from a stock (41 mM, 100% DMSO) was then added (2 mM final). The reaction was carried out at room temperature in the dark for one hour and then quenched with an excess of free L-cysteine. The resultant SapB^ANBD^ was syringe filtered through a PES 0.22 μm membrane and concentrated for a second size exclusion purification. Fractions aligning with the wild-type SapB elution volume peak were pooled and concentrated to determine the molarity and degree of labeling using the molar absorption coefficient of 22,500 cm^-1^M^-1^ and maximum wavelength of 478nm for the probe. The correction factor was determined by a parallel reaction of IANBD amide with BME and was determined to be 0.0396 (A280 / A478). The concentration of SapB^ANBD^ was calculated using the molar absorption coefficient of 3355 cm^-1^M^-1^ (Expasy, ProtParam) for SapB and the degree of labeling was approximately 0.73 moles of ANBD per mole of SapB. An 18-hour enzyme-activator assay was carried out incubating 4 μg of SapB^ANBD^ with and without GLA in the presence of Gb3-NBD and compared with the same assay using wild type SapB, as above.

### Chemical Cross-Linking Studies

*SapB titration*: SapB (0, 25, 50, 75, 100, 200 μM) and the catalytic nucleophile knockout GLA^D170A^ (10 μM) were incubated in the presence of BS^3^ (1 mM) in PBS pH 6.5. After one hour, the reaction was quenched with Tris pH 7.5 (26 mM final concentration) and 10 μL samples were run on a 12% acrylamide gel using Tricine-SDS-PAGE^80^ until the dye front reached the end of the gel. The gel was then stained with One-Step Blue (Biotium) for one hour and then de-stained in Mili-Q water overnight, after which they were imaged with a LI-COR Odyssey CLx and analyzed using the Image Studio Lite 5.2.5 software.

*Gb3-NBD titration*: SapB (50 μM) and GLA^D170A^ (10 μM) were incubated in the presence of BS^3^ (1 mM) and a range of Gb3-NBD concentrations (0, 50, 100, and 200 μM) in PBS pH 6.5 for one hour. Samples were processed as described above.

*BS*^3^ *titrations*– *Complex confirmation by NBD fluorescence and western blotting:* SapB (100 μM), SapB^ANBD^ (20 μM), GLA^D170A^ (10 μM), and Gb3-NBD (100 μM) were incubated in the presence of BS^3^ (0, 0.2, 0.5, 1 mM) in PBS pH 6.5 for one hour. SapB presence within the recognized SapB:GLA complex was confirmed by in-gel detection of SapB^ANBD^ (478 nm max absorbance) using an XcitaBlue transilluminator shield and GelGreen filter on a Gel Doc XR+ (BioRad) prior to One-Step Blue staining; GLA presence was confirmed by anti-GLA western blot of a duplicate gel, transferred with Towbin buffer to a PVDF membrane (Immobilon) using a semi-dry blotter (Cleaver Scientific) set to 38 V constant for 30 min. The membrane was blocked overnight with 5% milk in TBST and then serially incubated with polyclonal rabbit anti-GLA primary antibody N1C2 (Genetex) and secondary goat anti-rabbit IgG antibody (Rockland) diluted in TBST at room temperature for one hour for each incubation. The membrane was washed 3 times for 5 min in TBST between all steps. Finally, chemiluminescence detection of the blot was carried out using ECL Prime Western Blotting Kit (Amersham) and visualized on the Gel Doc XR+. *Controls*: To test saposin specificity, 100 μM SapD was used instead of SapB, and to test hydrolase specificity, 10 μM human acid-α-glucosidase (GAA) was used instead of GLA^D170A^ with samples analyzed as above. Finally, lower concentrations of SapB^ANBD^ (10 μM), GLA^D170A^ (1 μM), and Gb3-NBD (50 μM) were incubated in the presence of increasing concentrations of BS^3^ to test the cross-linking efficiency at protein concentrations near those used in the enzyme activation assay.

The *findSurfaceResidues.py* script in *PyMOL* (2.5 Å^2^ cutoff) was used to calculate the number of BS^3^ X-linker reactive, solvent accessible side chains on each protein monomer structure from the PDB (ID: 4V2O, SapB; 3BQQ, SapD; 3HG3, GLA; 5KZW, GAA).

This work is further detailed in the doctoral dissertation of corresponding author T.K.S. from ScholarWorks at the University of Massachusetts Amherst (Amherst, MA, USA).^81^

## Data availability

The atomic coordinates and structure factors for the SapB:Gb3-NBD and SapB:GLA complexes have been deposited to the Protein Data Bank under the accession codes 9AXG and 9AVS, respectively. Other data are available from the corresponding author upon reasonable request.

## ACKNOWLEDGMENTS

This research used resources of the Stanford Synchrotron Radiation Lightsource, SLAC National Accelerator Laboratory, which is supported by the U.S. Department of Energy, Office of Science, Office of Basic Energy Sciences under Contract No. DE-AC02-76SF00515. The SSRL Structural Molecular Biology Program is supported by the DOE Office of Biological and Environmental Research, and by the National Institutes of Health, National Institute of General Medical Sciences (P30GM133894). The contents of this publication are solely the responsibility of the authors and do not necessarily represent the official views of NIGMS or NIH. We thank Lisa Dunn, Clyde Smith, and Silvia Russi at SSRL. We thank the Biophysical and Mass Spectrometry Cores of the Institute for Applied Life Sciences - UMass Amherst (Amherst, MA, USA). This work was made possible by funding to S.C.G. from NIDDK at the National Institutes of Health, Award R01DK076877.

## AUTHOR CONTRIBUTIONS

T.K.S. designed research, performed experiments, and wrote the manuscript. S.C.G. designed and supervised research and wrote the manuscript. J.V.S provided guidance in chemical cross-linking assays and edited the manuscript. C.E.B. performed and analyzed the mass spectrometry of crystal samples and edited the manuscript. E.A. ran molecular dynamics simulations and edited the manuscript.

---

**Supplementary Figure 1:**
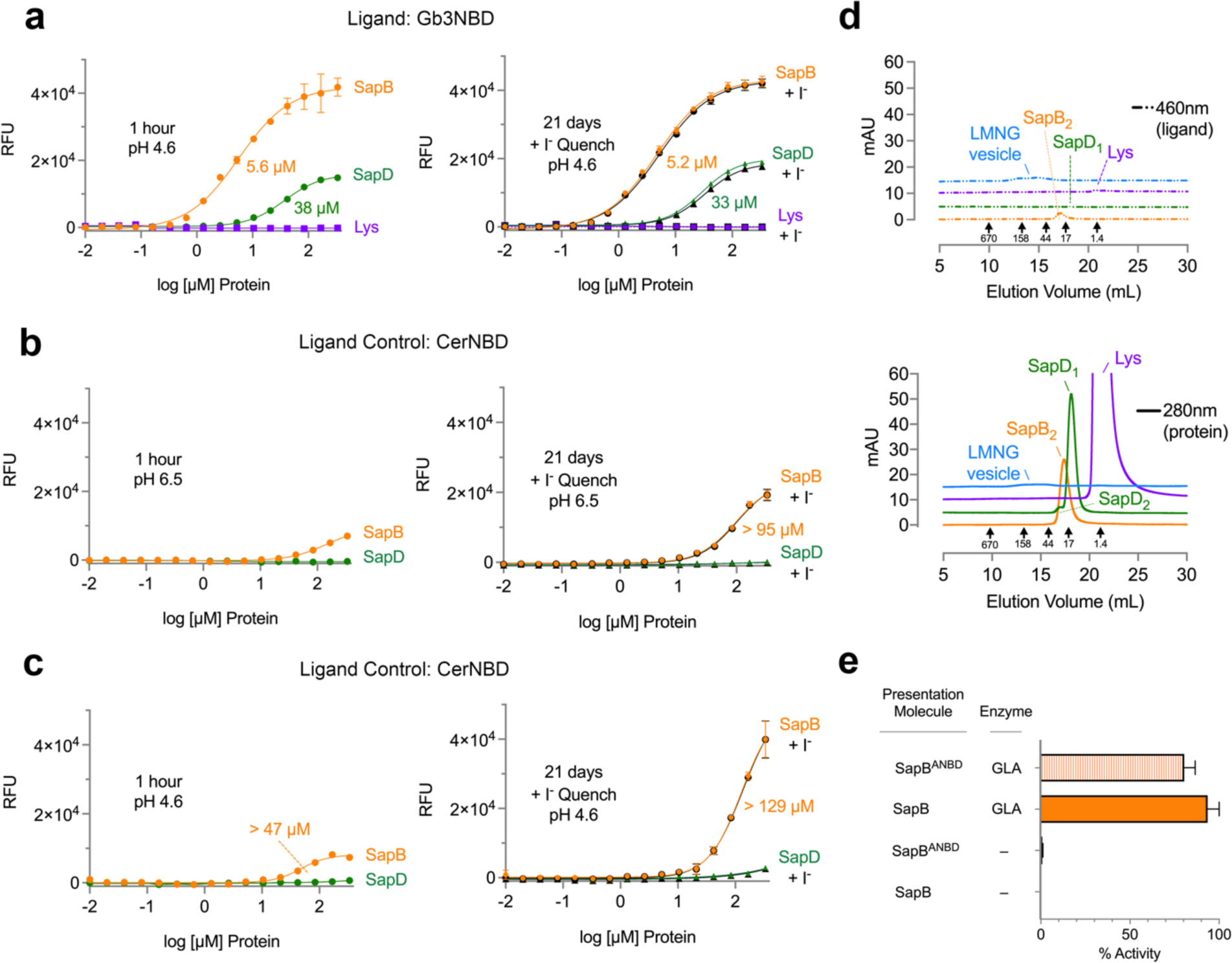
Specificity and ligand controls for SapB binding to Gb3-NBD. **a** The equilibrium binding of SapB (orange circle), the saposin specificity control SapD (green triangle), and negative control lysozyme (Lys, purple square) to Gb3-NBD after one hour (*left*) or after twenty-one days with iodide quenching (*right*) at pH 4.6. The apparent dissociation constants are labeled. **b** SapB and SapD binding to the ligand control Cer-NBD, which lacks the glycan of Gb3, after one hour (*left*) or twenty-one days later with iodide quenching (*right*) at pH 6.5. **c** The same as panel b but at pH 4.6. **d** The size exclusion chromatograms of SapB (orange), SapD (green), lysozyme (purple), and LMNG (blue) without Gb3-NBD ligand, in which dual wavelength monitoring tracks both Gb3-NBD ligand (top) and protein (bottom) absorbance. **e** Hydrolysis of Gb3-NBD, measured by an increase in the concentration of free galactose, using the fluorescent reporter SapB^ANBD^ (striped) or wild type SapB (orange) with and without wild type GLA. The concentration of free galactose detected was normalized to the highest value obtained by wild type SapB to show the comparative percent activity. The error bars in panels represent the standard deviation between replicates.

**Supplementary Figure 3:**
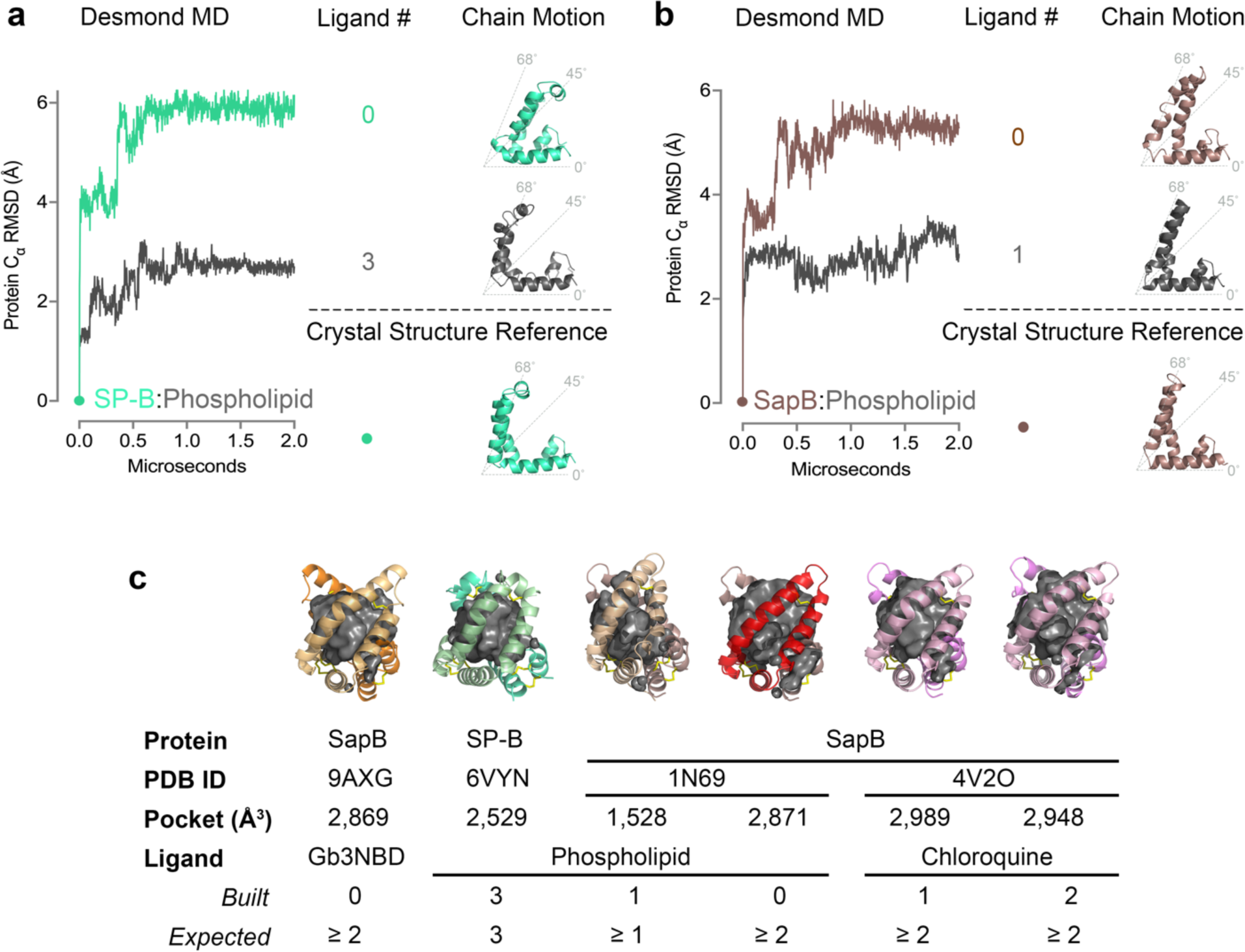
Molecular dynamics controls for interpreting the SapB:Gb3-NBD structure as ligand-bound. **a** Surfactant protein B (SP-B) bound to phospholipid (green-cyan) and **b** SapB bound to phospholipid (brown) dimers were used as positive controls for the MD simulations. The alpha carbon RMSD of the protein main chains is graphed by microsecond after the simulation. Representative intrachain motions as a function of ligand presence are shown to the right. **c** A comparative analysis of the ligand-bound states of the SapB:Gb3-NBD structure and controls, including the ‘unliganded’ dimer of SapB bound to phospholipid (red) and the dimers of SapB bound to chloroquine (pink). The calculated pocket volumes of the saposin and saposin-like dimers are shown for comparison. The number of cargos built within the dimers of the PDB structures is indicated and an estimate of expected ligand occupancy based on the pocket volume is provided, considering SP-B as a positive control.

**Supplementary Figure 4:**
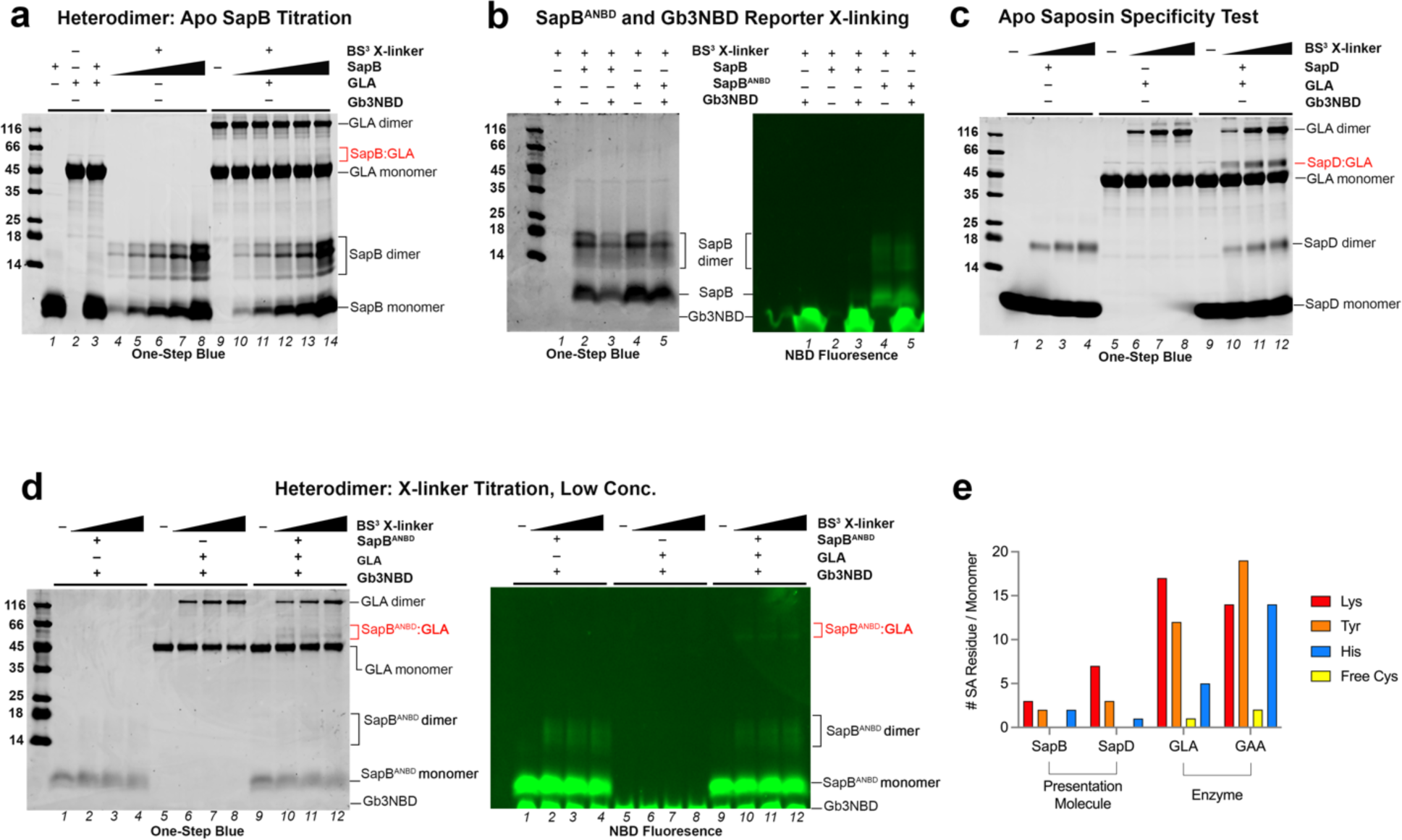
*In vitro* cross-linking controls to understand the specificity of the SapB:GLA interaction. **a** One-Step Blue stained SDS-PAGE gel of 10 μM GLA and increasing concentrations (0, 25, 50, 75, 100, and 200 μM) of SapB after incubation with 1 mM BS^3^ X-linker absent Gb3-NBD. **b** *Left*: SDS-PAGE of 100 μM SapB or SapB^ANBD^ after incubation with 1 mM BS^3^ X-linker in the presence and absence of 100 μM Gb3-NBD. *Right*: The NBD fluorescence of the SDS-PAGE gel prior to One-Step Blue staining to detect the NBD-containing bands using an XcitaBlue conversion screen and GelGreen detection filter**. c** SDS-PAGE gel of 10 μM GLA and 100 μM SapD samples after incubation with increasing concentrations of BS^3^ X-linker (0, 0.2, 0.5, 1mM) absent Gb3-NBD. **d** *Left*: SDS-PAGE of 1 μM GLA, 10 μM SapB^ANBD^, and 50 μM Gb3-NBD samples after incubation with increasing concentrations (0, 0.2, 0.5, 1 mM) of BS3 X-linker. *Right*: The NBD fluorescence of the SDS-PAGE gel prior to One-Step Blue staining. **e** A comparison graph of the number of BS^3^ X-linker reactive, solvent accessible (SA) side chains (Lys-red, Tyr-orange, His-blue, free Cys-yellow) on each protein monomer from PDB structures.

**Supplementary Figure 5:**
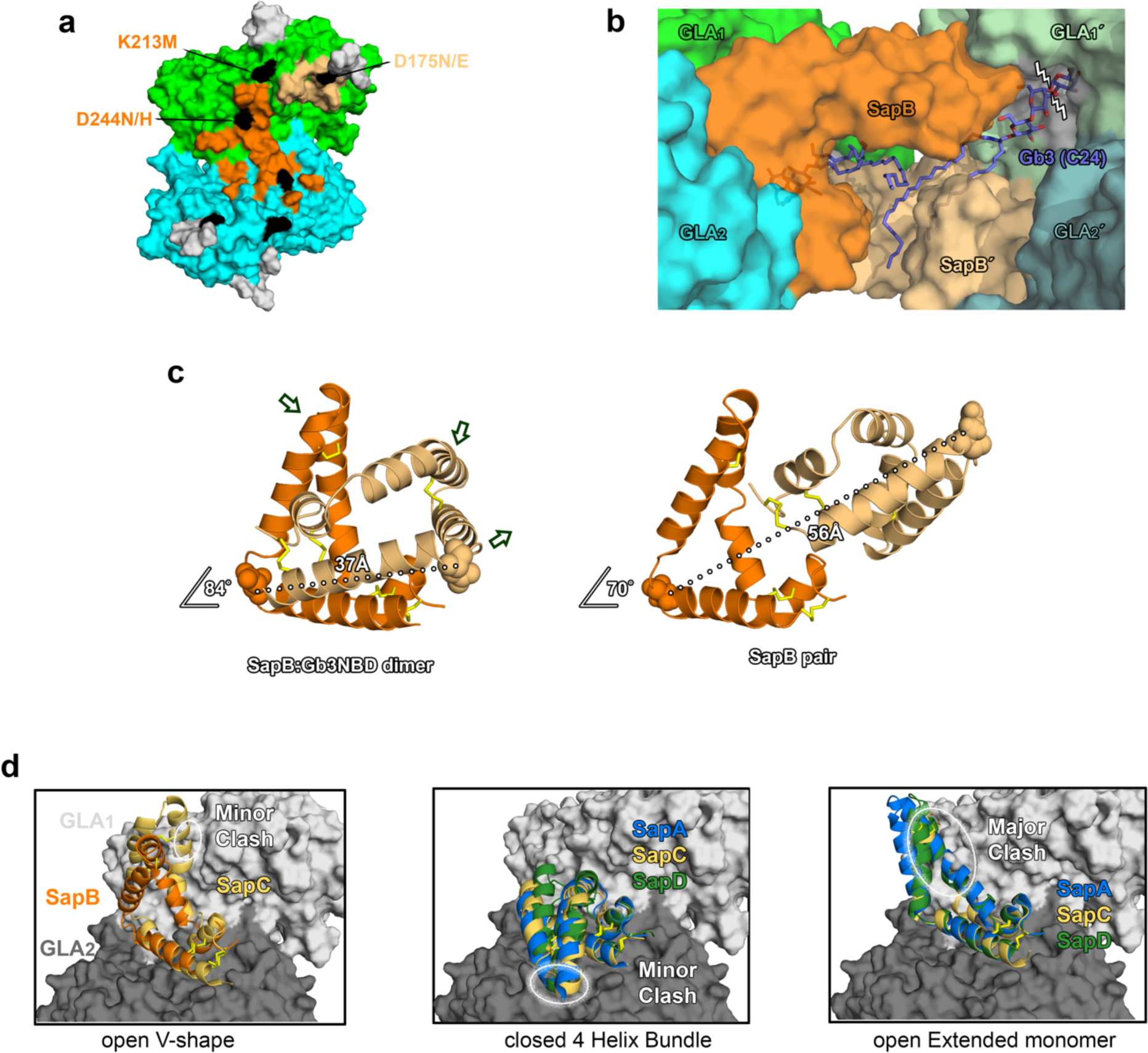
Models for the catabolism of Gb3 ligand by SapB:GLA and conformational importance. **a** Surface of the GLA dimer (green & cyan) with Fabry disease causing mutations (black) painted onto the SapB contact regions (orange & tan). Glycans are colored grey. **b** Surface of the SapB:GLA interaction with two Gb3 (C24) substrate molecules (purple sticks), manually docked according to torsion angle constraints. The structure of GLA dimer bound to melibiose was used to place the terminal αGal moiety in the GLA_1’_ (pale green) active site (grey) and guide the βGal exit. **c** Comparison between the SapB:Gb3-NBD dimer and the SapB crystallographic pair observed in the SapB:GLA binary complex. Arrows indicate the translations on the respective chains to move from the SapB dimer to the pair formation. The relative distances between the Asn21 Cβ (spheres) are shown for reference. **d** Alignments of SapA (blue), SapC (yellow) and SapD (green) crystal structures to the SapB (orange) chain making major contacts with the GLA dimer (light and dark grey). The models are organized by the saposin conformation and regions of major or minor clashes are indicated (white ovals).

**Supplementary Figure 6:**
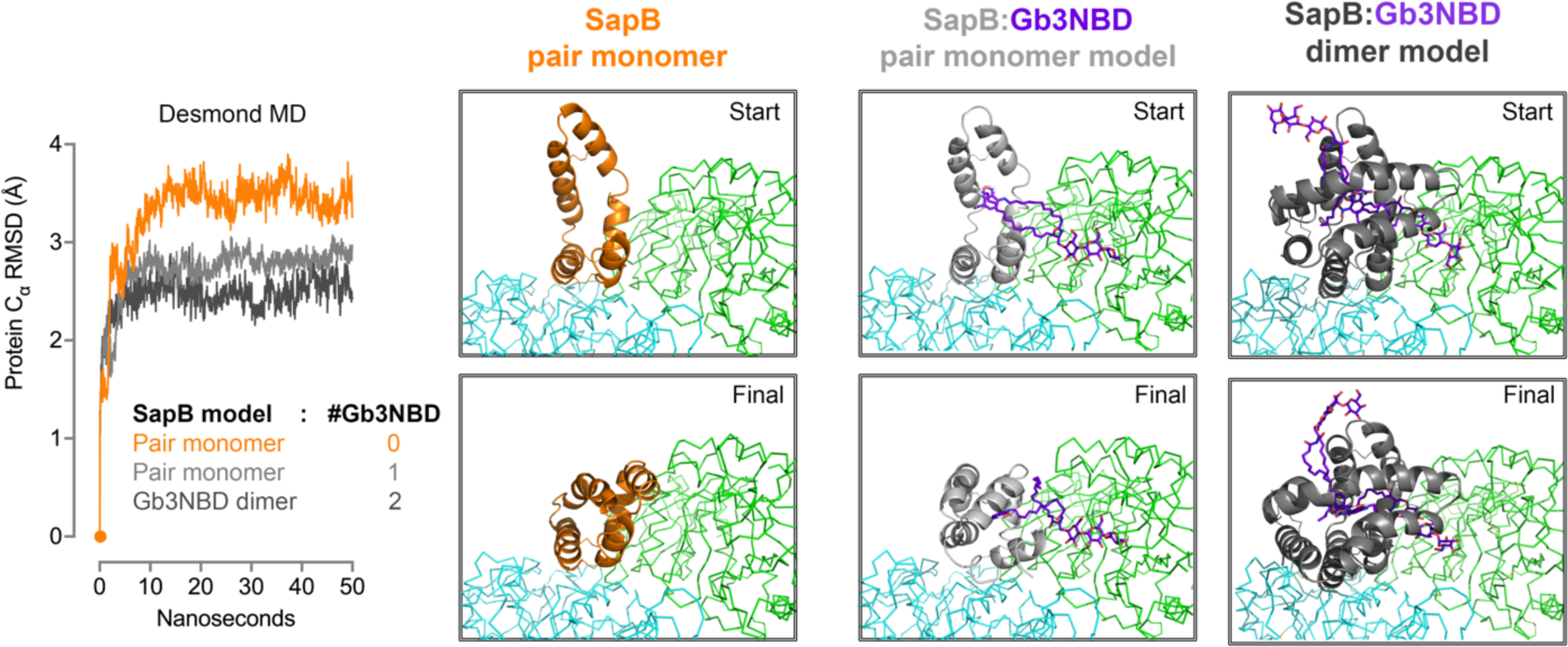
Molecular dynamics on the SapB:GLA binary complex provides models for substrate presentation. Simulations on the asymmetric unit components of the SapB:GLA crystals, containing a SapB monomer and GLA dimer, were conducted in the presence and absence of a single, modeled Gb3-NBD ligand before assessing the compatibility of a SapB dimer. *Left:* The total alpha carbon RMSD of the SapB and GLA chains is graphed by nanosecond after the MD simulation. The starting structure (orange dot) of each simulation is indicated. *Right:* Translations in the SapB chain (cartoon) from the start to the final state after 50 nanoseconds, both in the absence (orange) and presence (light grey) of a single Gb3-NBD molecule (purple stick), are shown. The GLA chain motions are shown in ribbon (green & cyan). A final simulation was rendered of a ligand-loaded SapB:Gb3-NBD dimer (dark grey), aligned to the SapB monomer in the crystal and slightly adjusted to avoid a minor hairpin clash into GLA.

